# Controlled and orthogonal partitioning of large particles into biomolecular condensates

**DOI:** 10.1101/2024.07.11.603072

**Authors:** Fleurie M. Kelley, Anas Ani, Emily G. Pinlac, Bridget Linders, Bruna Favetta, Mayur Barai, Yuchen Ma, Arjun Singh, Gregory L. Dignon, Yuwei Gu, Benjamin S. Schuster

## Abstract

Biomolecular condensates arising from liquid-liquid phase separation contribute to diverse cellular processes, such as gene expression. Partitioning of client molecules into condensates is critical to regulating the composition and function of condensates. Previous studies suggest that client size limits partitioning, with dextrans >5 nm excluded from condensates. Here, we asked whether larger particles, such as macromolecular complexes, can partition into condensates based on particle-condensate interactions. We sought to discover the biophysical principles that govern particle inclusion in or exclusion from condensates using polymer nanoparticles with tailored surface chemistries as models of macromolecular complexes. Particles coated with polyethylene glycol (PEG) did not partition into condensates. We next leveraged the PEGylated particles as an inert platform to which we conjugated specific adhesive moieties. Particles functionalized with biotin partitioned into condensates containing streptavidin, driven by high-affinity biotin-streptavidin binding. Oligonucleotide-decorated particles exhibited varying degrees of partitioning into condensates, depending on condensate composition. Partitioning of oligonucleotide-coated particles was tuned by altering salt concentration, oligonucleotide length, and oligonucleotide surface density. Remarkably, beads with distinct surface chemistries partitioned orthogonally into immiscible condensates. Based on our experiments, we conclude that arbitrarily large particles can controllably partition into biomolecular condensates given sufficiently strong condensate-particle interactions, a conclusion also supported by our coarse-grained molecular dynamics simulations and theory. These findings may provide insights into how various cellular processes are achieved based on partitioning of large clients into biomolecular condensates, as well as offer design principles for the development of drug delivery systems that selectively target disease-related biomolecular condensates.

**Significance Statement:** Biomolecular condensates are subcellular compartments that selectively recruit or exclude client molecules, even though condensates lack an enclosing membrane. Many biochemical reconstitution experiments have investigated mechanisms by which membraneless organelles control partitioning, modeling how cells spatiotemporally recruit components into condensates to regulate cellular functions. One outstanding question is whether partitioning is strictly limited by client size. In this work, we engineered nanoparticles with various sizes and surface functionalities and measured how these variables determine partitioning. We observed controlled and orthogonal partitioning of large particles into several condensate types, driven by strong particle-condensate interactions. Molecular simulations recapitulated key results. Our work advances understanding of how condensate composition is regulated, and our nanoparticle toolbox may also inspire a platform for drug delivery.

## Introduction

Condensation of biomolecules enables cells to form compartments without a surrounding membrane. Despite lack of a delimiting membrane, biomolecular condensates are chemically distinct from the cytoplasm and from one another, giving rise to unique condensate functions (1–3). A condensate’s biochemical components can be roughly divided into two categories: scaffolds (the biopolymers whose multivalent interactions drive phase separation and condensate formation) or clients (molecules that partition into condensates but which are not required for condensate assembly) (4). Significant effort has been devoted to understanding which clients partition into condensates, given the importance of this question to condensate biology. Most of these studies have focused on partitioning of small molecules, proteins, and nucleic acids. The view that has emerged is that small molecules may partition into condensates based on chemical compatibility; larger molecules, such as proteins and RNA, encounter additional barriers to partitioning, including that their presence in a condensate reduces the conformational entropy of scaffold biopolymers; but sufficient favorable interactions can permit protein and RNA clients to overcome this entropic penalty and partition into condensates (5–9).

This raises the question: Can even larger particles – on the order of tens or hundreds of nm – partition into condensates? Filling in this key knowledge gap is important for understanding how macromolecular assemblies, such as ribosomes, enzyme complexes, and viruses may partition into condensates. The limited existing data are mixed. Studies of dextran partitioning into biomolecular condensates *in vitro* and *in vivo* demonstrate a size-exclusion effect, where larger dextrans (especially > 70 kDa) tend to be excluded from condensates (10–12). On the other hand, recent studies suggest that the during HIV-1 infection, the intact virus capsid can transport through the nuclear pore complex, which is formed by condensation of FG repeat proteins (13, 14). An intermediate case is observed in P granules, where MEG-3 clusters adsorb to the condensate interface (15). A systematic investigation is required to test whether there is an upper limit to the size of clients that can partition into condensates, and to tease apart the biophysical principles that govern whether large clients partition into, are excluded from, or adsorb to the interface of condensates.

The central hypothesis of this study is that large, adhesive particles can partition into condensates via interactions with the condensate scaffold. We reason that this hypothesis will be valid if the condensate is terminally viscous, so that the biopolymer network can flow around the particles at time scales longer than the reptation relaxation time (16). To test this hypothesis, we engineered a toolbox of nanoparticles of varying sizes and surface chemistries, including particles that resist protein adhesion, particles that bind to scaffold proteins via specific protein-ligand interactions, and particles that interact non-specifically with condensates. We studied the partitioning of these particles into three model *in vitro* condensates: the intrinsically disordered RGG domain from LAF-1 (17, 18), the SARS-CoV-2 nucleocapsid (N) protein (19–21), and a cationic artificial intrinsically disordered protein (22). These experiments, along with coarse-grained molecular dynamics simulations, allowed us to investigate the interplay between particle size and stickiness on partitioning into condensates. Our studies indicate that arbitrarily large particles can partition into condensates based on adhesive interactions between the particle and condensate, but larger particles require greater adhesion strength to do so. Particles that do not interact with condensates are excluded; as the interaction strength increases, the particles localize to the interface of condensates; and as the interaction strength increases yet further, the particles partition into the condensates. Remarkably, large particle partitioning can be highly tunable and specific, allowing orthogonal partitioning in which two different particle types can target two immiscible condensates. Together, this work addresses the fundamental biophysical question of how size and stickiness determine client partitioning into condensates, which is critical for understanding how condensates regulate their composition in the complex cellular milieu, and it may inform how clients can be engineered to partition into condensates for therapeutic intervention.

## Results

### Partitioning into condensates depends on client size and binding interactions

We first examined the permeability of condensates to various sizes of dextran: 10 kDa, 40 kDa, and 70 kDa, corresponding to hydrodynamic radii of 1.9, 4.8, and 6.5 nm, respectively (23). We mixed rhodamine-labeled dextrans with three representative condensate-forming proteins: the LAF-1 RGG domain; the SARS-CoV-2 nucleocapsid (N) protein, which phase separates when mixed with RNA; and a designed polycationic repeat polypeptide, (GRGNSPYS)_25_ (SI Appendix, Fig. S1) (22). We observed that partitioning into condensates is inversely related to dextran size, consistent with previous studies (Fig. 1A) (10, 24). The partition coefficient of 70 kDa dextran was 5-fold lower than that of 10 kDa dextran for all three condensates, and it dropped below 1 for LAF-1 RGG and N protein. These results confirm that size plays an important role in client partitioning into biomolecular condensates, as established in the literature, and suggest that the condensates have a mesh size of very roughly 5 nm (10).

**Fig. 1.**
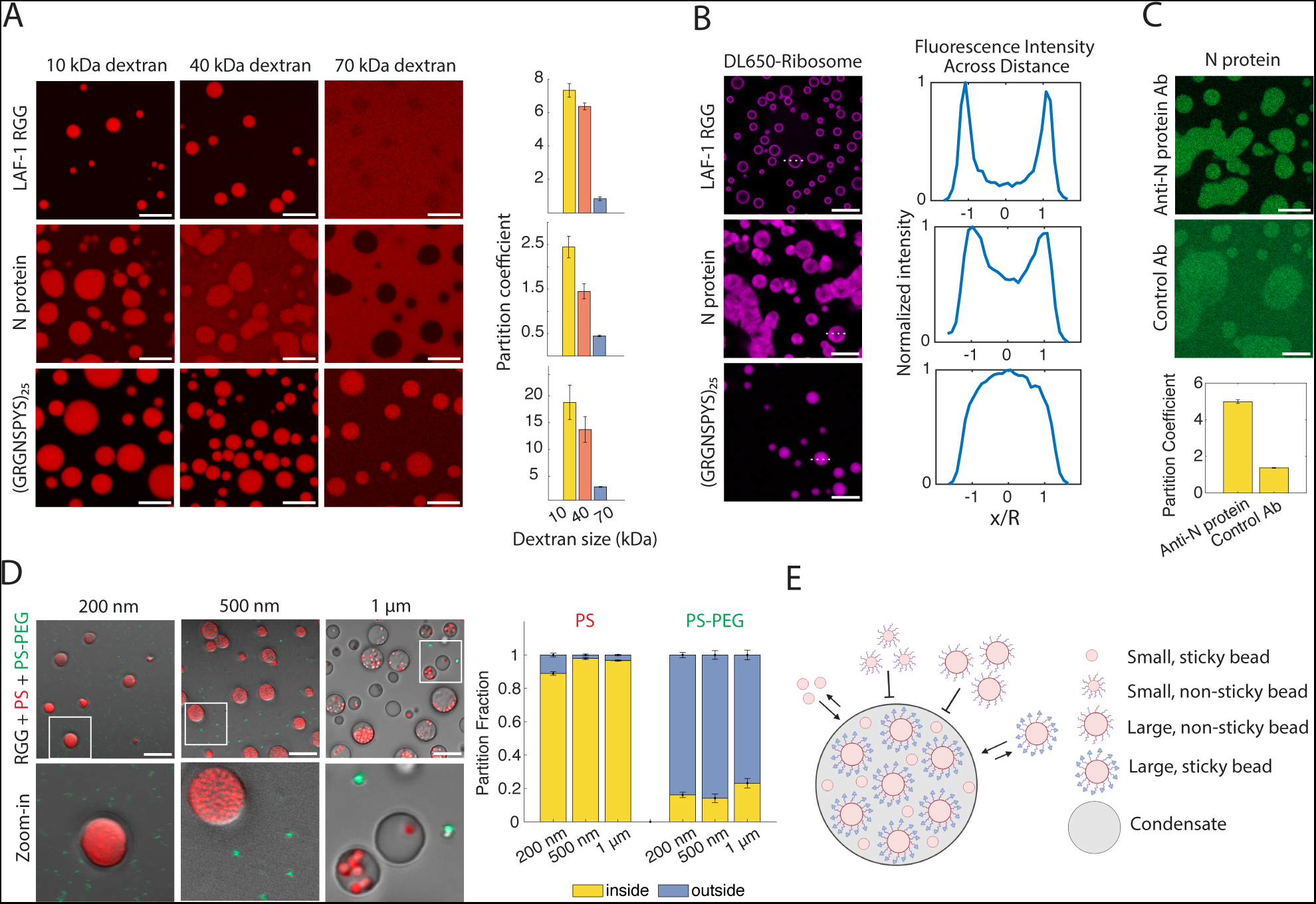
Condensate permeability and partitioning characterized by a variety of probes. (A) 10 kDa, 40 kDa, and 70 kDa rhodamine-labeled dextrans were mixed with LAF-1 RGG, (GRGNSPYS)_25_, or N protein. For all figures, buffer was 150 mM NaCl 20 mM Tris, pH 7.5, unless stated otherwise. (Right) Partition coefficients were quantified for each protein and each dextran size. (B) *E. coli* ribosomes, labeled with DyLight 650 (DL650), were mixed with each protein. Buffer included 1 mM MgCl_2_. (Right) Fluorescence intensity line profiles are shown for representative condensates (corresponding to dashed white lines on micrographs). (C) N protein was mixed with anti-N protein IgG or isotype control antibody. (Below) Partition coefficients were quantified for each antibody. (D) RGG was mixed with PS (red) and PS-PEG (green) beads of various sizes. PS beads as large as 1μm partition into RGG condensates. PS-PEG beads are excluded from condensates. (Right) Partition fractions were quantified for each bead type. Any particles inside the condensate or at its interface were counted as inside, and all other particles were counted as outside. Error bars represent SEM with *n* = 20. For all figures, n equals number of images from at least 2 independent experiments, unless otherwise stated. (E) Schematic illustrating particle partitioning into condensates. Unmodified PS beads partition into condensates, while PEGylated beads are excluded. Hypothesis: adding sticky moieties, depicted as blue triangles, to PEGylated beads may recruit beads back into condensates. Scale bars, 10 μm.

However, hydrophilic and uncharged dextrans are expected to interact only weakly with condensates. We hypothesized that for other clients, interactions with the scaffold protein can drive partitioning into condensates of clients > 5 nm. We therefore examined whether larger macromolecular assemblies can partition into condensates. Several studies have hypothesized that actively-translating cellular puncta called translation factories are biomolecular condensates (25–27). This raises the question of whether ribosomes can partition into biomolecular condensates. To test this, we fluorescently labeled *E. coli* ribosomes, which are about 21 nm in diameter, and measured their partition coefficient into condensates (Fig. 1B and SI Appendix, Fig. S2). Ribosomes bound to the periphery of LAF-1 RGG condensates, partitioned heterogeneously into N protein condensates, and partitioned uniformly into GRGNSPYS droplets. These differences can be rationalized by the fact that all three proteins are cationic, but LAF-1 RGG is the least so (net charge ≈ +3 at neutral pH), while GRGNSPYS is highly cationic (net charge ≈ +25), and N protein is known to be a promiscuous RNA-binding protein (net charge ≈ +23). These observations suggest that particles significantly larger than the dextrans may partition into condensates, depending on condensate-particle interaction strength.

We next sought to directly test the hypothesis that strong binding interactions between condensate scaffold proteins and clients can drive partitioning, even for large clients. To explore this, we compared partitioning into N protein condensates of antibodies with and without specific affinity for N protein. The approximate molecular weight of IgG is 150 kDa, and its dimensions are about 14.5 nm x 8.5 nm x 4.0 nm (28). Based on our dextran experiments, 70 kDa dextran was mostly excluded from N protein condensates, with a partition coefficient of < 0.5. In contrast, despite its larger size, anti-N protein IgG was enriched in N protein condensates, with a partition coefficient of about 5 (Fig. 1C). Isotype control antibody showed only weak partitioning (partition coefficient of about 1.5, suggesting that control antibodies have more non-specific interactions with condensates than dextrans). These results further demonstrate that, based on affinity, molecules larger than would be expected from the dextran experiments can partition into condensates.

Intrigued by the ribosome and antibody partitioning results, we asked whether even larger particles can partition into condensates. We mixed carboxyl-modified polystyrene beads (PS) of various sizes with LAF-1 RGG protein. Beads even up to 1 µm diameter partitioned into the condensates (Fig. 1D), agreeing with prior microrheology studies (29–31). The PS bead surface is negatively charged due to the carboxylate moieties but retains hydrophobic character from the polystyrene, so multiple non-specific interactions likely contribute to the partitioning of the PS beads into condensates.

Prior studies have demonstrated that coating nanoparticles with a dense brush of low molecular-weight polyethylene glycol (PEG) reduces protein adsorption to the particles (32, 33). Therefore, we hypothesized that PEGylating beads will cause the beads to be excluded from condensates. To test this, we conjugated 5 kDa PEG to the PS beads to form PS-PEG particles (the PS beads are negatively charged, whereas the PS-PEG beads have near-neutral zeta-potential; SI Appendix Table 1). Strikingly, the PS-PEG beads (200 nm, 500 nm, and 1 µm diameter) were essentially completely excluded from the LAF-1 RGG condensates (Fig. 1D). Based on the dichotomy between PS and PS-PEG bead partitioning, we conclude that sticky beads of arbitrary size can partition into condensates, whereas non-sticky beads are excluded. Our findings clearly indicate that sticky interactions can significantly facilitate the partitioning of particles into condensates. Although the PS beads partition due to nonspecific interactions, our results so far suggest that functionalizing particles with biomolecules that have specific interactions with condensates may enable targeted partitioning of clients into these condensates (Fig. 1E).

### Simulations demonstrate competing effects of size and protein-particle attraction

To further elucidate the effects observed from experiments, we conducted molecular dynamics simulations of a simple binary system of Lennard-Jones (LJ) particles. Employing this simplified model allows for us to access more significant size disparities between component types than would be otherwise allowed by more complex coarse-grained simulation models. We thus simulate a condensed phase of “protein” particles at conditions where phase separation is observed, i.e. below the critical temperature and above the saturation concentration. We then add “bead” particles to the system to represent large cargo molecules such as dextrans, antibodies, or ribosomes. Although we can simulate significant size disparities between different particle types, we still cannot achieve disparities as large as those observed with the protein-polystyrene bead experiments, as those would require a prohibitively large number of protein particles in the simulation. For all cases, we fixed the total volume fraction ratio of protein:bead to 5:1 to be consistent with conditions tested in experiments.

Using this system, we tested the effects of both size and stickiness of the beads on partitioning into a condensate by varying two LJ parameters, namely the diameter of the beads, σ_2_, and the protein-bead interaction energy, ε_12_ (see SI Methods). We first studied the effect of size on partitioning by looking at two cases where beads and protein are placed together in a phase-separating system. In Fig. 2A, we show the density profile of protein, which forms a dense phase at the center of the box, and a dilute phase outside. Keeping the protein-bead interactions constant at ε_12_ = 0.8 (i.e. 80% of protein-protein interaction strength), we find that when beads are the same size as the protein particles (σ_2_ = σ_1_ = 1.0), they are enriched in the dense phase, but when the diameter of the beads is increased to σ_2_ = 2.0 (now 8x the volume of the protein), they are completely excluded from the dense phase. This is analogous to the case of the 10 and 70 kDa dextrans, where the smaller client partitions into the condensate, and the larger is excluded.

**Fig. 2.**
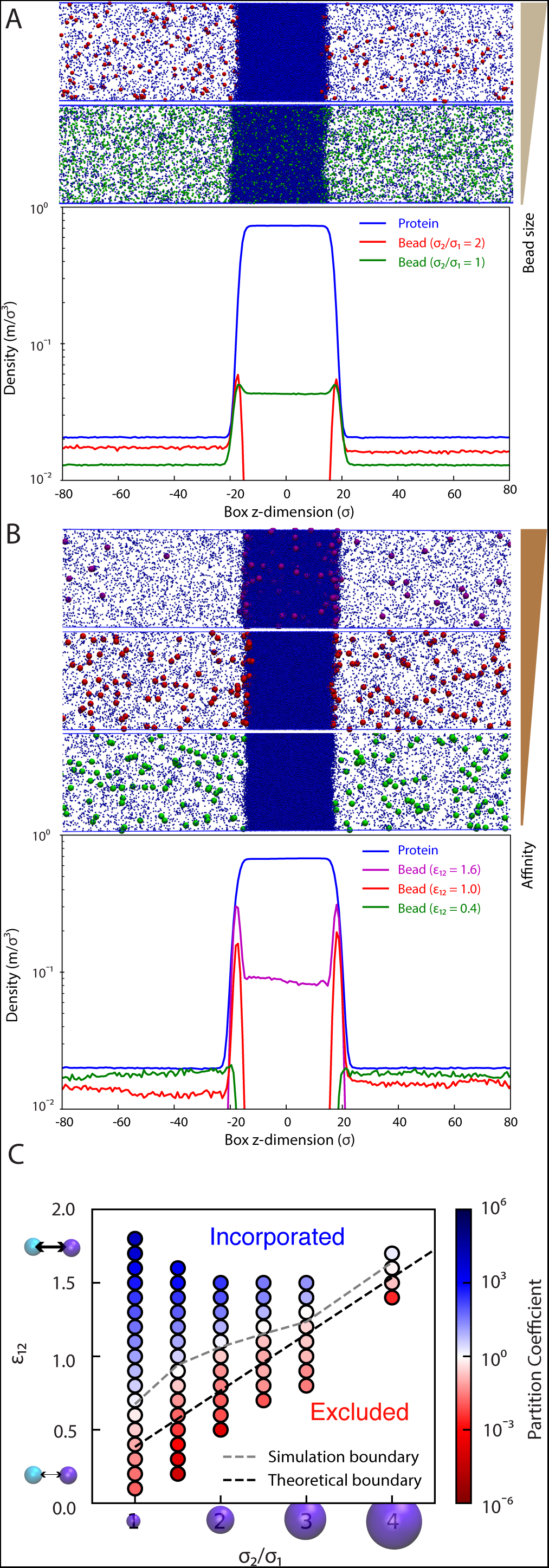
Simulations demonstrate effect of size disparity and interaction energies on partitioning. (A) Density profile of “protein” and “bead” components in slab simulations of Lennard-Jones particles demonstrating the effect of bead size on partitioning. Smaller beads incorporate most easily, while larger beads are excluded. Inset shows snapshots from simulations. (B) Density profile of “protein” component and “bead” component in slab simulations, showing that stronger protein-bead interactions result in incorporation of beads into the condensate, while weaker protein-bead interactions result in exclusion of beads. Inset shows snapshots from simulation. (C) A grid of simulation parameters was tested, demonstrating that partition coefficients increase with decreasing size and with increasing protein-bead interaction energy. The dashed black line shows the crossover point where partition coefficient is equal to 1 according to theory derived in Eq. 3, and the dashed gray line shows the crossover point from the simulations.

The effect of increased size opposing particle partitioning into condensates can be offset, however, by increasing the attractive interactions between the protein particles and beads. In Fig. 2B, we show that for a system of particles of size σ_2_ = 3.0 (27x the volume of protein beads; a disparity similar to ribosome and protein sizes), weak attractive interactions result in full exclusion, while strong attractive interactions result in preferential incorporation. We also find that at intermediate interaction strengths, the beads localize to the surface of the dense phase, but do not partition inside. Similar surface enrichment has been observed previously for proteins and protein clusters at condensate interfaces (15, 18, 34). Thus, we largely recapitulate the observations from experiments showing that ribosomes and other large clients such as nanoparticles can be either preferentially incorporated or excluded from condensates, or localized to the surface, and that varying protein-client interaction strength is sufficient to capture each of the differential localizations.

We then asked how size and protein-bead attraction compete to control the partitioning of beads into a protein-rich dense phase. In Fig. 2C, we show a grid of conditions tested with varying both σ_2_ and ε_12,_ finding that smaller beads and beads with stronger attractive interactions are more preferentially incorporated into the dense phase, while larger beads and those with weaker attractive interactions are preferentially excluded from the dense phase.

To explain the reason for each of these effects within the simplified LJ model, we formulate an analytical expression (see SI Appendix) to represent the energetic component of the transfer free energy (i.e. change in energy of the system upon insertion of a single particle into a dense phase of protein):

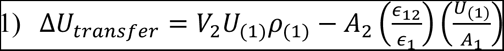

where V_2_ and A_2_ are the volume and surface area of a bead; U_(1)_ and ρ_(1)_ are the average energy per protein particle and average number density, respectively, within a pure condensate of protein particles; and A_1_ is the surface area of a protein particle. The first term describes the unfavorable energetic penalty of displacing protein particles within a certain volume in the condensed phase having an energy density of U_(1)_ρ_(1)_. The second term describes the favorable interactions formed between the inserted bead and the condensate-forming protein particles. Expanding the volume and area terms for a spherical bead, we obtain:

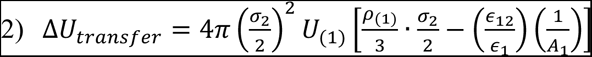

By solving for ΔU_transfer_ = 0, we can obtain a linear function to describe the boundary between preferential incorporation and exclusion, assuming negligible entropic contribution to the transfer free energy. This gives a linear relationship between ε_12_ and σ_2_ and depends only on the surface area and the condensed phase density of the protein beads:

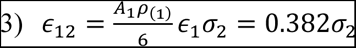

The density we use in this calculation was obtained by using a case where beads were not incorporated into the condensate (i.e. ε_12_ < 0.5) and fitting the dense phase of protein, yielding ρ_(1)_ = 0.73. We have also used the fact that in all the simulations, *∈*_1_ = 1 and σ_1_ = 1. This boundary between incorporation and exclusion is shown as a dashed line in Fig. 2C and shows reasonable agreement with the simulation data, particularly for larger bead sizes. Possible sources of error include the larger concentration of beads in simulations (elaborated in SI Text).

From this section, we conclude that an arbitrarily large bead can partition into a condensate provided that the bead’s interactions with protein can be made arbitrarily large. We also present evidence that the increase of stickiness needs to be proportional to the radius (not volume) of the particles. Finally, we note that the simulations also suggest an explanation for enrichment of particles at the condensate surface. Since the protein density ρ_(1)_ decreases along the axis normal to the condensate surface, the energy penalty (first term in Eq. 1) from adding a large particle decreases at the interface, resulting in an enrichment of beads at the interface.

### Partitioning can be controlled through specific binding

Based on the simulations and experimental results so far, we hypothesized that diverse protein-ligand interactions can overcome the thermodynamic penalty that would otherwise exclude large particles from condensates. One of the strongest known non-covalent protein-ligand interactions is between streptavidin and biotin (35). The pair has an unusually high affinity of K_d_ ~ 10^−15^ M as well as high specificity, and we therefore sought to harness this interaction to control particle partitioning (Fig. 3A). To test this, we fused streptavidin to the RGG domain to create a streptavidin-RGG fusion protein (SA-RGG) (SI Appendix, Fig. S1). To prepare the particles, we started from the premise that PS beads must first be PEGylated to block non-specific interactions, with biotin then displayed at the free end of the PEG. We therefore prepared PS-PEG-biotin beads. Remarkably, 90% of PS-PEG-biotin beads with 500 nm diameter partitioned into SA-RGG condensates, whereas 85% of the control particles (500 nm PS-PEG) were excluded (Fig. 3B and SI Appendix, Fig. S3). The result was the same whether the beads were added before or after the condensates formed (SI Appendix, Fig. S4). These results indicate that specific, high-affinity interactions can drive the thermodynamic partitioning of large particles into biomolecular condensates.

**Fig. 3.**
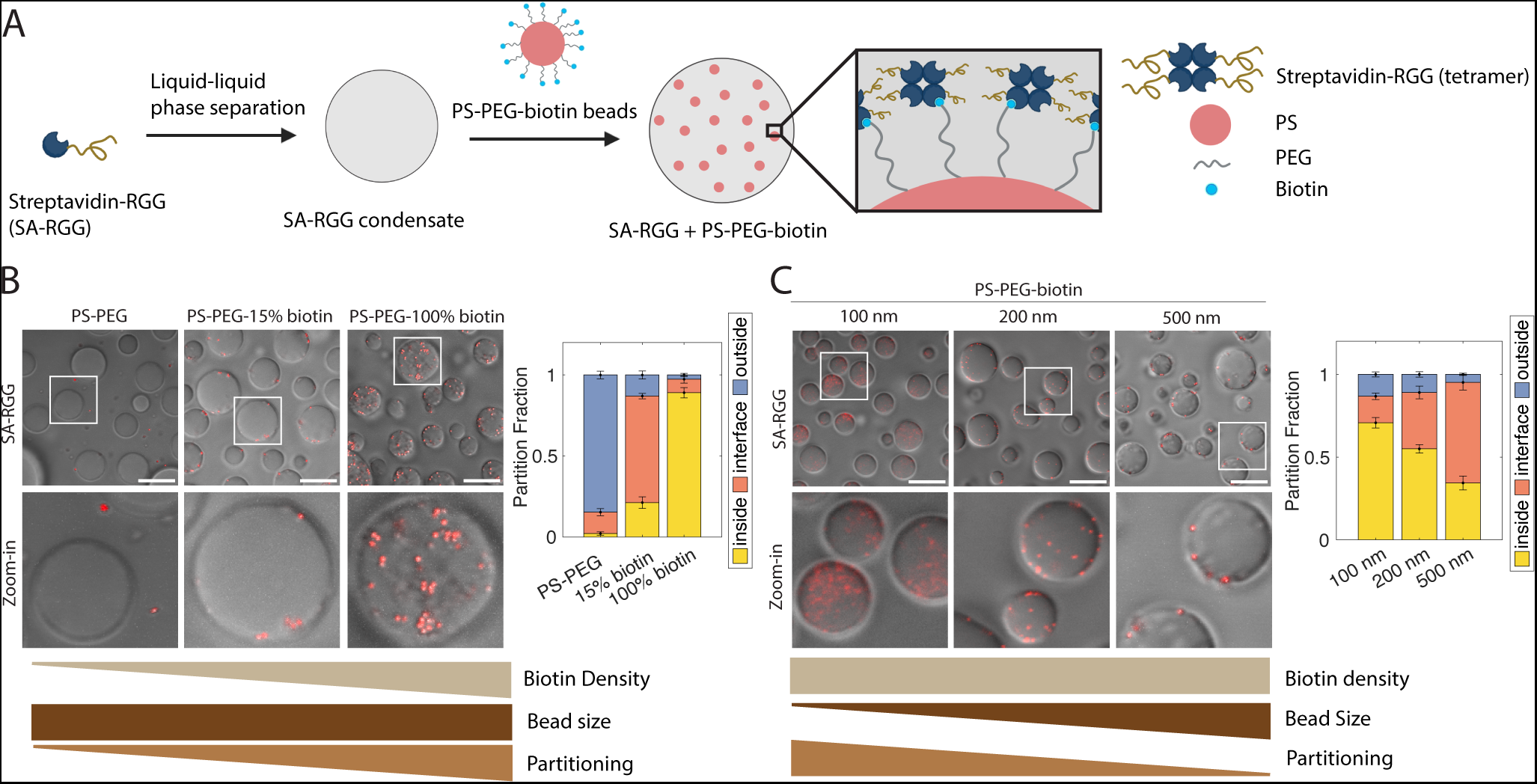
Streptavidin-biotin interactions drive bead partitioning into condensates. (A) Schematic model of PS-PEG-biotin partitioning into SA-RGG condensates. Streptavidin (SA, which tetramerizes) binds biotin, resulting in bead recruitment into condensates. (B) Increasing biotin surface density on particles leads to increased partitioning of beads into SA-RGG. (Right) Partition fractions were quantified for each bead. (C) When biotin surface density is held constant, smaller beads display higher partitioning. (Right) Partition fractions were quantified for each bead. Scale bars, 10 μm. Error bars represent SEM with *n* = 10.

Interestingly, when the PS-PEG-biotin particles were prepared (SI Methods) with reduced biotin surface density (15%), the particles predominantly adsorbed to the SA-RGG condensate interface (Fig. 3B). This agrees with the simulation results, in which beads of intermediate interaction strength localized at the condensate periphery rather than inside or outside the condensates.

To further explore the interplay of size and affinity, we prepared various PS-PEG-biotin particle sizes and analyzed partitioning into SA-RGG condensates. To avoid confounding variables, the biotin surface density was kept consistent across the different sized particles, at an intermediate biotin density (0.02 biotin/nm^2^). We observed that the partition fraction decreased with increasing bead size (Fig. 3C): 100 nm beads predominantly partitioned into the condensates, whereas 200 nm beads were partially inside and partially at the interface, and 500 nm beads were predominantly localized to the condensate interface. This result demonstrates that larger size indeed hinders partitioning of particles into condensates, given consistent ligand area density on the particles, which is also in agreement with the simulations at constant surface interaction strengths.

### Protein-nucleic acid interactions can drive particle partitioning

Our experiments with ribosomes (Fig. 1B) suggest that protein-nucleic acid interactions can drive particle partitioning into condensates. We therefore asked whether these interactions can be harnessed to engineer particles for controlled partitioning. To test this, we again began with PS-PEG particles, which resist protein adhesion and are excluded from condensates (SI Appendix, Fig. S5). We further modified the PS-PEG beads by conjugating DNA oligonucleotides to the free end of the PEG using strain-promoted azide-alkye cycloaddition click chemistry (Fig. 4A). We first studied polyA20 conjugated to 500 nm PS-PEG (PS-PEG-polyA20). We observed that these PS-PEG-polyA20 beads partitioned robustly into N protein and GRGNSPYS condensates but adsorbed to the surface of LAF-1 RGG condensates (Fig. 4B), all in 150 mM NaCl buffer. Interestingly, the PS-PEG-polyA20 bead partitioning displayed protein sequence dependence that qualitatively agrees with the trends we observed for ribosome partitioning, presumably because both are determined by condensate-nucleic acid interactions.

**Fig. 4.**
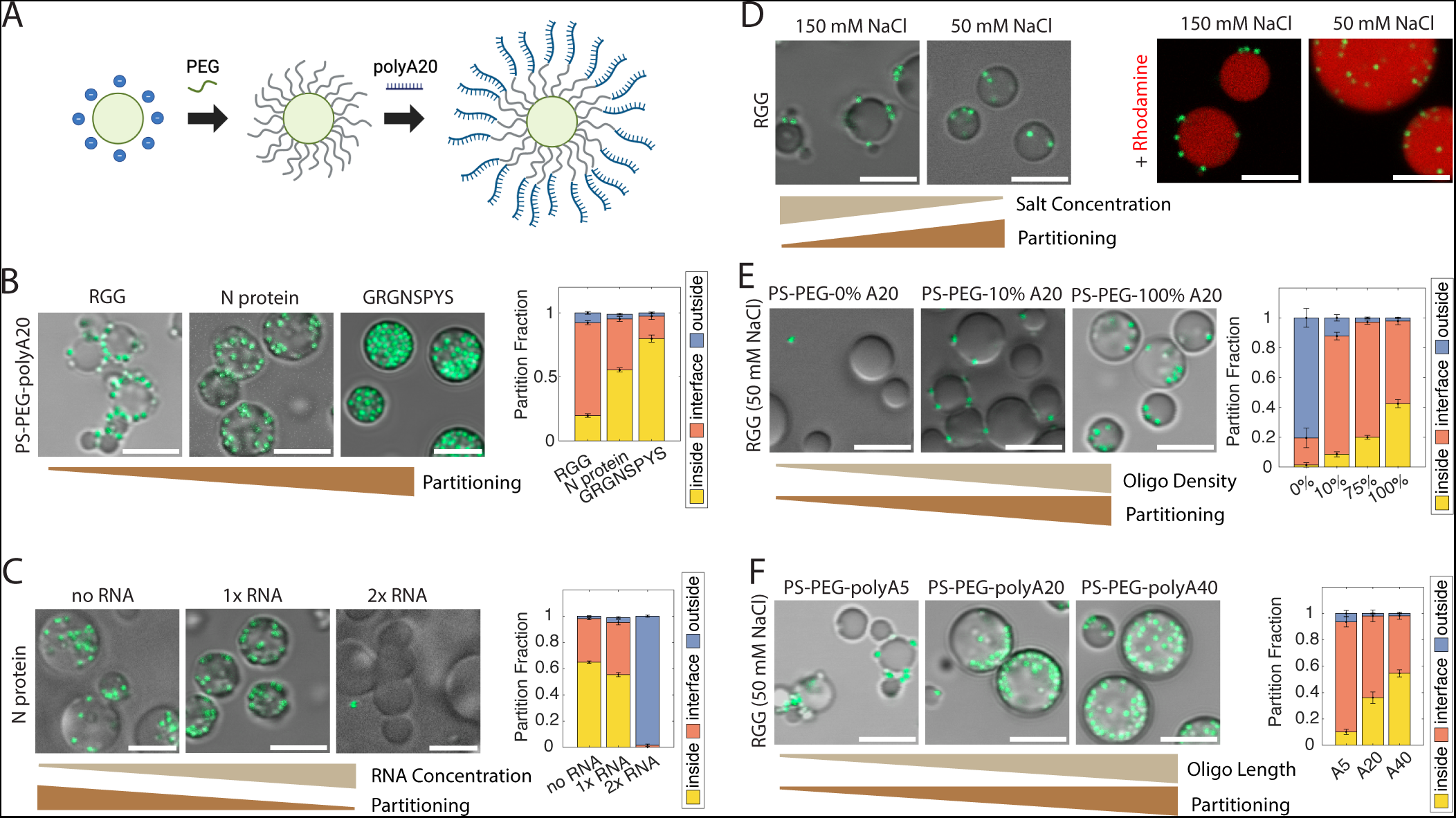
PS-PEG-oligonucleotide beads partition into condensates based on electrostatic interactions. (A) Schematic of PS particle surface functionalization to PS-PEG and then PS-PEG-polyA20. (B) Proteins were mixed with 500 nm PS-PEG-polyA20 (green) beads. Particles mainly stick to the interface of RGG condensates, but partition more robustly into N protein and GRGNSPYS condensates. (Right) Partition fractions were quantified for each sample. (C) Increasing RNA concentration leads to decreased partitioning of PS-PEG-polyA20 into N protein condensates. (Right) Bead partition fractions were quantified at each RNA concentration. (Scale bars, 5 μm.) (D) 500 nm PS-PEG-polyA20 beads partition into RGG more at lower salt concentration, and less at higher salt. (Right) Rhodamine was added to better visualize particle location with respect to condensate interface. (E) Partitioning of 500 nm beads with 0, 10, and 100% oligo density. Increased oligonucleotide surface density on beads leads to greater partitioning into condensates. (Right) Partition fractions were quantified for 0, 10, 75, and 100% oligo density. (F) PS-PEG-polyA5, A20, and A40 partitioning into RGG condensates at 50 mM NaCl. Longer oligonucleotides leads to higher partitioning into condensates. (Right) Partition fractions were quantified for each bead. Error bars represent SEM with *n* ≥ 10.

Previously (Figs. 1A,B and 4B), N protein was mixed with polyA RNA to induce its phase separation. We hypothesized that there could be competing effects between PS-PEG-polyA and polyA RNA for binding to N protein. To test this, we compared the partition coefficient of PS-PEG-polyA20 particles into N protein condensates with varying polyA RNA concentrations: no RNA, 1x RNA (0.5 mg/mL, the RNA concentration used in our prior N protein experiments), or 2x RNA (1 mg/mL). In the case of no RNA, 8 kDa PEG was used as a crowding agent to induce phase separation of N protein. Absence of RNA led to more robust partitioning of PS-PEG-polyA beads into condensates, whereas 2x RNA concentration resulted in >98% of PS-PEG-polyA beads being excluded from the condensates (Fig. 4C). This result demonstrates that “free” nucleic acids can compete with particle-conjugated nucleic acids for binding to N protein, and hints at a possible mechanism by which condensates exclude unwanted clients in cells.

Since electrostatic attraction between polyA20 and condensates likely plays an important role in PS-PEG-polyA20 partitioning, we hypothesized that varying salt concentration will alter partitioning, as cations in solution will screen the negatively charged oligonucleotides. To test this, we compared partitioning of the beads at different NaCl concentrations (Fig. 4D). Whereas most PS-PEG-polyA20 beads adsorbed at the interface of RGG condensates in 150 mM NaCl buffer, the beads partitioned further into the condensates in 50 mM NaCl buffer. Upon fluorescently staining the condensates, we observed that at 150 mM NaCl, the interfacial beads are partially immersed in the RGG phase and partially immersed in the dilute aqueous phase, whereas at 50 mM NaCl, even beads near the interface are entirely or almost entirely immersed in the RGG phase (our resolution is insufficient for accurate contact angle measurements). Similarly, PS-PEG-polyA20 beads partitioned robustly into GRGNSPYS condensates at lower salt concentration, whereas they remained localized at the condensate interface at higher salt concentration (SI Appendix, Fig. S6). We conclude that electrostatic interactions play an important role in driving partitioning of PS-PEG-polyA20 beads into these condensates.

We conducted additional experiments to verify our results with PS-PEG-polyA20 beads. Orthogonal projections and three-dimensional renderings from confocal microscopy confirm inclusion of PS-PEG-polyA20 and exclusion of control beads from condensates (SI Appendix, Fig. S7). We tested beads of a different size (200 nm PS-PEG-polyA20), and separately, we tested beads with a different oligonucleotide sequence (PS-PEG-polyT20). Both partitioned into condensates, to different degrees (SI Appendix, Fig. S8). Beads of various sizes conjugated with polyA20 hybridized to polyT20 oligonucleotides (PS-PEG-polyA20/T20) also partitioned into condensates (SI Appendix, Fig. S9). Overall, these results demonstrate that large particles coated with single or double-stranded oligonucleotides can partition into biomolecular condensates, provided that the condensate-nucleic acid interactions are sufficiently strong.

To further develop this idea, we next asked whether density of oligonucleotide on the particle surface tunes partitioning. We prepared beads functionalized with varying surface densities of polyA20 (0%, 10%, 75%, and 100%; see SI Methods). We mixed these beads with LAF-1 RGG in 50 mM NaCl buffer and observed that increasing oligonucleotide surface density increased partitioning (Fig. 4E and SI Appendix, Fig. S10). Similarly, we asked whether particle partitioning could be tuned based on oligonucleotide length. We therefore compared PS-PEG-polyA beads with different polyA lengths: PS-PEG-polyA5, PS-PEG-polyA20, and PS-PEG-polyA40. We tested the partitioning of these beads into RGG in 50 mM NaCl buffer. Indeed, increasing oligonucleotide length resulted in increased partitioning (Fig. 4F). PS-PEG-polyA5 beads were predominantly localized at the condensate interface, whereas nearly 40% of PS-PEG-polyA20 beads and 60% of PS-PEG-polyA40 beads partitioned into the condensates. Together, these results suggest that the strength of condensate-nucleic acid interaction determines PS-PEG-polyA particle partitioning and can be tuned by polyA attachment density or length.

### Reversing particle partitioning

So far, we have demonstrated how nanoparticle surface chemistry can be engineered to drive particle partitioning into condensates. Next, we asked whether condensate-particle interactions can be blocked to prevent partitioning or to expel partitioned particles. Building on Fig. 4C, we sought to understand whether competition between free and particle-bound ligand depends on which species binds to the condensate first. When we mixed LAF-1 RGG condensates (at 50 mM NaCl) with free Cy5-labeled polyA20 (i.e. DNA strands that are not attached to particles), and then subsequently added PS-PEG-polyA20 beads, we observed that the oligos partitioned into the condensates whereas the beads were excluded (adsorbed to the condensate interface) – both when observed after 10 min and after 24 hr (Fig. 5A). Similarly, when SA-RGG condensates were mixed with free fluorescently labeled biotin (biotin-4-fluorescein) before adding in PS-PEG-biotin beads, the free biotin partitioned but the beads were excluded – both when observed after 10 min or 24 hr. Thus, in both systems, adding abundant free ligand first can occupy binding sites and prevent partitioning of beads. However, a difference was observed when particles were added first, before the free ligand (Fig. 5B). When we first allowed the PS-PEG-polyA20 beads to partition into RGG condensates before adding free Cy5-polyA20, after 10 minutes of equilibration the beads remained partitioned inside even while the oligos also partitioned in, but after 24 hr, the beads were predominantly excluded (adsorbed to the interface) of similarly sized droplets. In contrast, PS-PEG-biotin beads that had partitioned into SA-RGG condensates remained partitioned even after subsequent addition of biotin-4-fluorescein – both when observed after 10 min and after 24 hr. We hypothesize that the equilibrium partitioning is achieved for PS-PEG-polyA20 beads within 24 hours, resulting in free ligand displacing the beads, whereas the PS-PEG-biotin beads are kinetically trapped due to the long lifetime of biotin-streptavidin bonds (36).

**Fig. 5.**
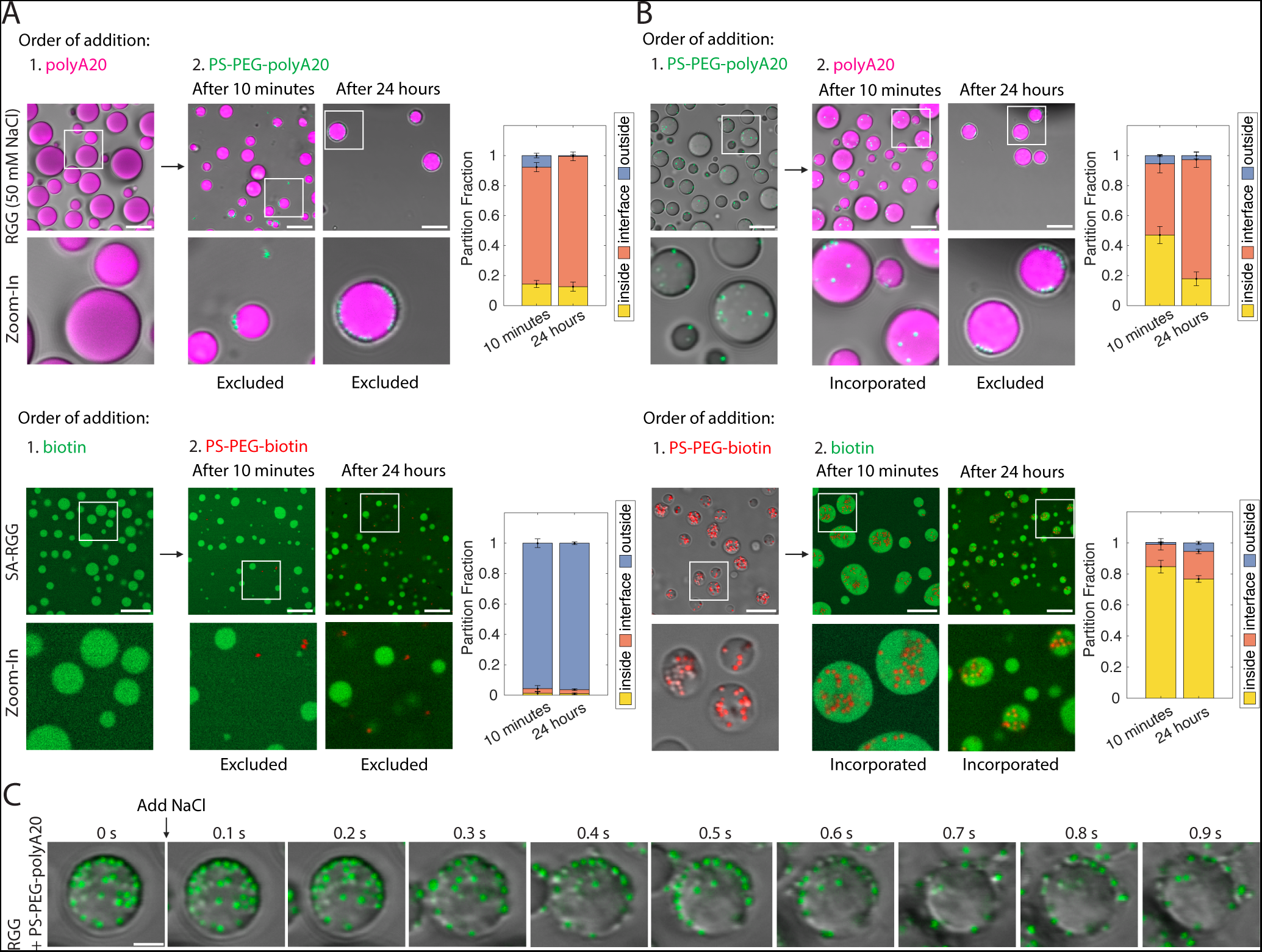
Expulsion of beads from condensates. (A) Free ligands prevent beads from partitioning into condensates. (Top) Free Cy5-polyA20 prevents PS-PEG-polyA20 (green) from partitioning into RGG condensates. (Bottom) Biotin-4-fluorescein prevents PS-PEG-biotin (red) from partitioning into SA-RGG. (B) Kinetic trapping of beads in condensates. (Top) When Cy5-polyA20 is added after PS-PEG-polyA20 already partitioned in, beads stay inside condensates after 10 minutes, but are excluded after 24 hr equilibration. (Bottom) Adding biotin-4-fluorescein after PS-PEG-biotin already partitioned in results in beads remaining inside condensates, even at 24 hr. (Scale bars, 10 μm.) (Right) Partition fractions were quantified. Error bars represent SEM with *n* ≥ 8. (C) Time lapse images of beads being expelled from condensates. RGG with PS-PEG-polyA20 (green) was originally at 50 mM NaCl. Salt concentration was raised to 150 mM NaCl at t = 0. Beads rapidly transport out of the condensates. (Scale bar, 5 μm.)

This motivated us to ask whether directly weakening the particle-condensate interaction could expel partitioned beads. We have seen that PS-PEG-polyA20 partitioning is salt concentration dependent (Fig. 4D), so we hypothesized that once the beads are partitioned in, raising the salt concentration may expel the beads from the condensates. We conducted this experiment with PS-PEG-polyA20 particles partitioned into RGG condensates at 50 mM NaCl. Raising the salt concentration from 50 mM to 150 mM NaCl triggered immediate and rapid transport of PS-PEG-polyA20 outward from the condensates (Fig. 5C). After equilibration, the sample (now at 150 mM NaCl) displayed the same droplet morphology and particle partitioning as samples freshly prepared with the same salt concentration (SI Appendix, Fig. S11 compared to Fig. 4D). This result suggests that increasing the salt concentration has a rapid and drastic effect on the energetic landscape inside the condensates.

### Orthogonal particle partitioning

We have established that distinct protein-ligand interactions can drive particles to partition into various condensates, and that the strength of these interactions are tunable. We were curious whether these features permit selective partitioning. We first asked whether the interface and interior of condensates could be simultaneously targeted by tuning interaction strength. Indeed, when we mixed both PS-PEG-polyA5 and PS-PEG-polyA40 beads with LAF-1 RGG, we observed that PS-PEG-polyA40 beads predominantly partitioned into the condensates, while PS-PEG-polyA5 adsorbed to the interface (Fig. 6A). Similarly, PS-PEG-100%biotin beads partitioned into SA-RGG condensates while simultaneously PS-PEG-10%biotin was localized at the interface, due to the beads’ high (100%) vs. low (10%) biotin surface density. This difference is highlighted by a radial density profile of the beads (Fig. 6A). Similar results were observed when RGG was mixed with both PS-PEG-10%polyA20 and PS-PEG-100%polyA20 (SI Appendix, Fig. S12).

**Fig. 6.**
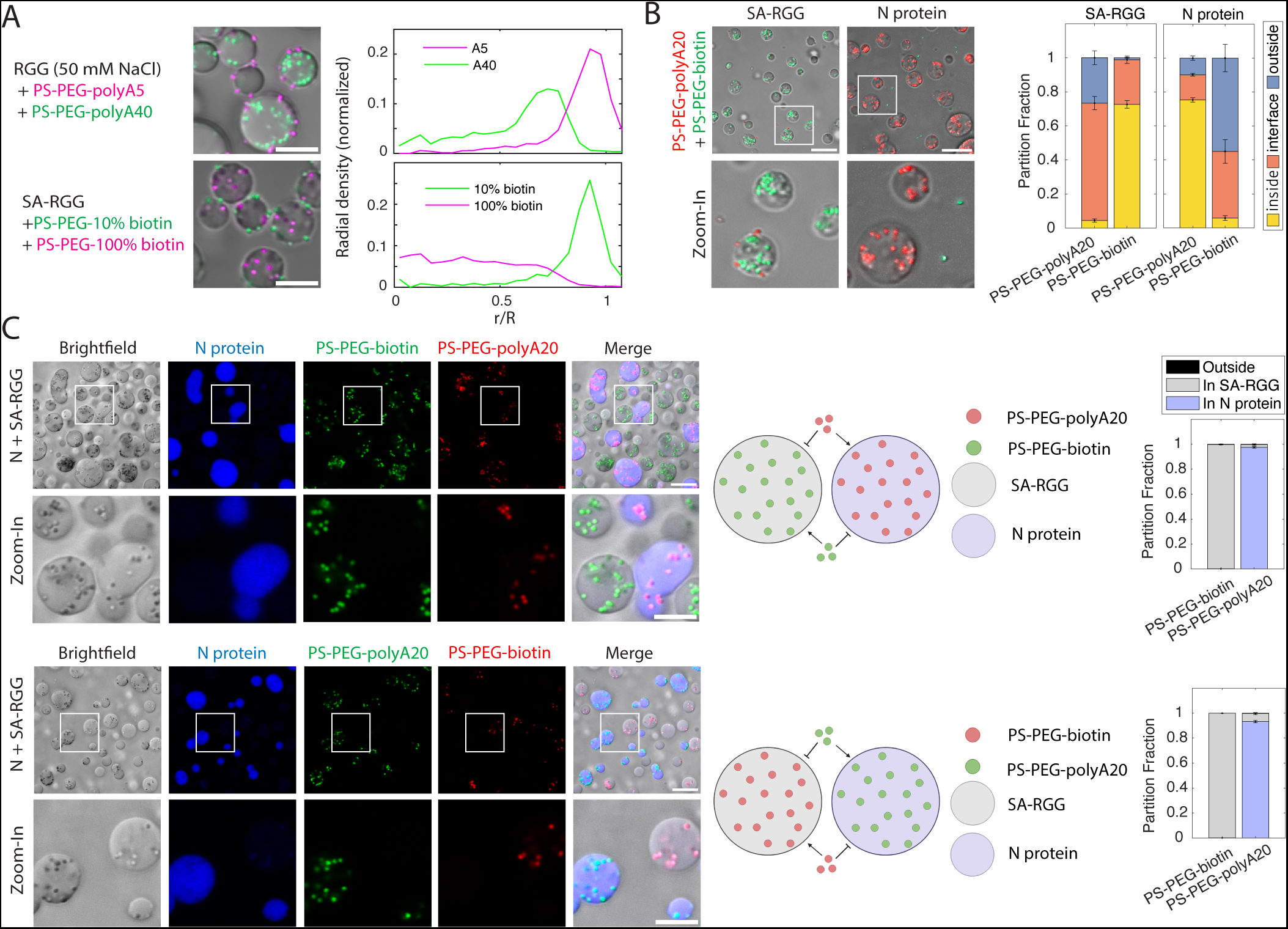
Orthogonality of bead partitioning into condensates. (A) Selective partitioning based on condensate-particle interaction strength. PS-PEG-polyA5 (magenta) adsorbs to condensate interface, while PS-PEG-polyA40 (green) partitions into RGG condensates. Likewise, PS-PEG-10% biotin (green) adsorbs to condensate interface, while PS-PEG-100% biotin (magenta) partitions. (Right) Radial density profile shows bead concentration as a function of normalized distance from condensate center. (B) Orthogonality of PS-PEG-biotin and PS-PEG-polyA20 partitioning into SA-RGG and N protein condensates. PS-PEG-biotin (green) partitions into SA-RGG, but not into N protein. PS-PEG-polyA20 (red) partitions into N protein, but not SA-RGG condensates. (Right) Partition fractions were quantified for each bead. (C) When SA-RGG is mixed with AF647-labeled N protein, they phase separate into immiscible condensates. PS-PEG-polyA20 only partitions into N protein condensates, while PS-PEG-biotin only partitions into SA-RGG. (Middle) Schematic of PS-PEG-polyA20 partitioning into AF647-labeled N protein and PS-PEG-biotin partitioning into unlabeled SA-RGG. (Right) Partition fraction quantified. Error bars represent SEM with *n* ≥ 10. (Scale bars, 10 μm.)

We next tested how beads with distinct surface chemistries would partition when combined. We therefore mixed PS-PEG-polyA20 and PS-PEG-biotin particles with SA-RGG, and separately, with N protein. In line with our previous results, PS-PEG-biotin partitioned robustly into SA-RGG condensates (in 150 mM NaCl buffer), while PS-PEG-polyA20 remained at the periphery (Fig. 6B). Conversely, PS-PEG-polyA20 partitioned robustly into N protein condensates, while PS-PEG-biotin beads were excluded. We conclude that distinct surface chemistries can target particles to distinct condensate microenvironments.

Building upon this result, we asked whether orthogonality of bead-condensate interactions can be observed when SA-RGG and N protein condensates are combined. To test this, we mixed SA-RGG with labeled N protein and added both green PS-PEG-biotin and red PS-PEG-polyA20 beads. Strikingly, SA-RGG and N protein formed immiscible and distinct condensates. PS-PEG-biotin beads partitioned overwhelmingly (>99%) into the SA-RGG condensates, whereas PS-PEG-polyA20 partitioned overwhelmingly (>97%) into the N protein condensates (Fig. 6C). Consistent results were obtained with green PS-PEG-polyA20 beads and red PS-PEG-biotin beads, confirming that the orthogonal partitioning was due to which biomolecules are decorated on the particle surface, and not due to any property of the particle core, such as interactions involving the fluorescent dyes. These results demonstrate that large particles can orthogonally target distinct condensates in a multi-component system, based on selective particle-condensate interactions.

## Discussion

In this paper, we hypothesized that strong client-scaffold interactions can enable large particles to overcome thermodynamic barriers and partition into liquid condensates. To test this hypothesis, we engineered a toolbox of nanoparticles with diverse sizes and surface chemistries. We found that PEG-coated beads resist partitioning into condensates. In contrast, biotin-functionalized beads partition into streptavidin-tagged condensates based on the high-affinity interaction between biotin and streptavidin, and oligonucleotide-conjugated beads can partition into condensates via protein-nucleic acid interactions – even for beads as large as 1 µm. Partitioning was tuned by modifying oligonucleotide length and surface density, or biotin density, thereby modifying the interaction strength between bead and condensate.

This work expands our understanding of “who’s in and who’s out” of biomolecular condensates in two significant ways (4). One key biophysical insight of our experiments and theory is that arbitrarily large particles can partition into liquid condensates, provided that condensate-particle interactions are sufficiently strong. Based on the large size of the beads in our experiments and their stickiness, we reason that the particles diffuse not by hopping between entanglement cages, but rather by waiting for condensate polymers to rearrange and flow at timescales longer than the relaxation time (16, 37, 38). Particle-protein interactions may also contribute to network rearrangement. A second key insight is that large particles functionalized with orthogonal surface chemistries can selectively target immiscible condensates. Previous studies have shown that small molecules with favorable physicochemical properties partition into condensates (5). Previous studies have also demonstrated that larger (~5 nm), “inert” molecules are excluded from condensates, whereas proteins and nucleic acids that interact with condensates can partition (39). However, it was unknown whether larger particles can be controllably and orthogonally targeted to biomolecular condensates. Our work demonstrates that the answer is yes, with important implications for biology and bioengineering.

With respect to condensate biology, our study may provide insights into how partitioning is spatiotemporally regulated in cellular condensates. Many large biomolecular complexes are believed to assemble and/or function in condensates (e.g., ribosomes, RNA PolII, spliceosomes) (40, 41), and our study quantifies the tradeoff between size, adhesiveness, and partitioning of such complexes. For instance, our work is consistent with the proposed mechanism of vectorial flux of ribosomal precursors in the nucleolus (42). In this model, relatively nascent rRNA transcripts interact more strongly with scaffold components (such as NPM1 and SURF6) in the granular component of the nucleolus. However, as the transcripts mature and bind to ribosomal proteins, the number of multivalent binding sites for scaffold proteins is reduced, thereby thermodynamically driving the expulsion of fully assembled pre-ribosomal particles from the nucleolus. Our work is also consistent with the finding that large cargo, such as the HIV-1 capsid, may transport through the nuclear pore complex (NPC) based on adhesive interactions with FG domains that, by competition, “melt” inter-FG-repeat interactions (13, 14). In synthetic biology, it has been demonstrated that synthetic organelles can be hubs for specific and selective protein translation (43–45). Our study implies that a key to the design of new condensates for orthogonal protein translation is engineering robust recruitment of ribosomes into the synthetic condensates.

Besides those tested here, many other small molecule ligands and biopolymers could be used to target particles into condensates. For instance, a logical extension is to explore oligonucleotide sequence-specific partitioning. The diversity of possible particle functionalities presents several opportunities. One is the design of probes for microrheology measurements of condensates. It should be possible to design particles to partition into any predominantly viscous biomolecular condensate, and additionally, different functionalities could be used to test the effect of probe surface chemistry on condensate rheology. Although our experiments were conducted *in vitro*, future studies may explore whether surface-engineered particles can partition into condensates *in vivo*, similar to how previous studies measured the partitioning of microinjected dextrans into *in vivo* condensates, such as nucleoli, NPCs, and P granules (11, 12). Microinjected or cell-penetrating particles could perhaps then be used to measure condensate rheology *in vivo* as well.

Our work also suggests exciting possibilities for drug delivery. Some of the main challenges in drug delivery are limited penetration to the targeted microenvironment (46) and nonspecific distribution (47). To address these challenges, modification of nanoparticle size and surface properties, such as PEGylation and tissue targeting moieties, have been extensively explored in nanoparticle-based drug delivery (48). Our work suggests that similar approaches can be extended to drug delivery to cellular condensates. Our experiments demonstrated orthogonal, specific, and efficient partitioning of particles into condensates due to avidity. This suggests it may be possible to engineer particles to target disease-related biomolecular condensates, such as in neurodegenerative diseases (49), cancer (50, 51), and viral infection (52).

There are numerous biomolecular condensates in cells, having different compositions and carrying out various crucial roles. How do these condensates regulate what goes in and out? How can we leverage these properties for biotechnology applications? Our work suggests there is no size limit to partitioning and demonstrates orthogonal partitioning of particles into condensates. The principles revealed in this paper can serve as a foundation to help elucidate how biological condensates regulate partitioning and can be harnessed for therapeutically targeting condensates.

## Methods

Refer to SI Appendix for details. All genes of interest were cloned into bacterial expression vectors in frame with 6x-His tags. Proteins were recombinantly expressed in *E. coli* and purified by affinity chromatography. Beads were PEGylated by carboxyl-to-amine coupling following a previously described protocol (33). Oligonucleotides were synthesized by solid-phase synthesis and then conjugated to PEGylated beads using click chemistry, as previously described (53). Confocal and brightfield microscopy were used to measure partitioning. MD simulations were conducted using the LAMMPS simulation engine (54).

## Acknowledgements

We thank Cristobal Garcia Garcia, Kristi L. Kiick, Shiv Rekhi, and Jeetain Mittal for designing and providing the GRGNSPYS protein. We also thank Xinyi Li for purifying GRGNSPYS protein. We thank Richard Haber for use of the Zetasizer. We gratefully acknowledge Elizabeth Nance for the PEGylation protocol. We thank Nina Shapley, Ned Wingreen, and Yaojun Zhang for helpful discussions. We also thank Dr. Srinivas Chakravartula for help with oligonucleotide characterization. This work was supported by NIH grants R35GM142903 (to B.S.) and R35GM150589 (to G.D.). This work was also supported by Rutgers University startup funds (to Y.G. and G.D.) and computing resources through Rutgers Engineering Computing Services. F.K. was supported by NIH training grant T32 GM135141 and B.L. was supported by NSF REU award 2149971.

## Author contributions

Fleurie M. Kelley: Experimental design, data collection and analysis, writing—original draft, writing—review and editing. Anas Ani: Data collection and analysis, writing—review and editing. Emily G. Pinlac: Data collection and analysis, writing—review and editing. Bridget Linders: Data collection. Bruna Favetta: Provision of protein, feedback on manuscript. Mayur Barai: Provision of protein, feedback on manuscript. Yuchen Ma: Provision of oligonucleotides. Arjun Singh: Software development, data analysis. Gregory L. Dignon: Supervision, software development, data analysis, writing—original draft, writing—review and editing, funding acquisition. Yuwei Gu: Supervision, experimental design, provision of oligonucleotides, writing—original draft, writing—review and editing, funding acquisition. Benjamin S. Schuster: Conceptualization, supervision, experimental design, software development, data analysis, writing—original draft, writing—review and editing, funding acquisition.

## References

1. C. P. Brangwynne, et al., Germline P Granules Are Liquid Droplets That Localize by Controlled Dissolution/Condensation. Science 324, 1729–1732 (2009).

2. S. F. Banani, H. O. Lee, A. A. Hyman, M. K. Rosen, Biomolecular condensates: organizers of cellular biochemistry. Nat. Rev. Mol. Cell Biol. 18, 285–298 (2017).

3. C. P. Brangwynne, T. J. Mitchison, A. A. Hyman, Active liquid-like behavior of nucleoli determines their size and shape in Xenopus laevis oocytes. Proc. Natl. Acad. Sci. 108, 4334–4339 (2011).

4. J. A. Ditlev, L. B. Case, M. K. Rosen, Who’s In and Who’s Out—Compositional Control of Biomolecular Condensates. J. Mol. Biol. 430, 4666–4684 (2018).

5. S. A. Thody, et al., Small Molecule Properties Define Partitioning into Biomolecular Condensates. [Preprint] (2022). Available at: https://www.biorxiv.org/content/10.1101/2022.12.19.521099v1 [Accessed 24 June 2024].

6. H. R. Kilgore, et al., Distinct chemical environments in biomolecular condensates. Nat. Chem. Biol. 20, 291–301 (2024).

7. T. J. Nott, T. D. Craggs, A. J. Baldwin, Membraneless organelles can melt nucleic acid duplexes and act as biomolecular filters. Nat. Chem. 8, 569–575 (2016).

8. S. Choi, M. O. Meyer, P. C. Bevilacqua, C. D. Keating, Phase-specific RNA accumulation and duplex thermodynamics in multiphase coacervate models for membraneless organelles. Nat. Chem. 14, 1110–1117 (2022).

9. D. M. Mitrea, M. Mittasch, B. F. Gomes, I. A. Klein, M. A. Murcko, Modulating biomolecular condensates: a novel approach to drug discovery. Nat. Rev. Drug Discov. 21, 841–862 (2022).

10. M.-T. Wei, et al., Phase behaviour of disordered proteins underlying low density and high permeability of liquid organelles. Nat. Chem. 9, 1118–1125 (2017).

11. D. L. Updike, S. J. Hachey, J. Kreher, S. Strome, P granules extend the nuclear pore complex environment in the C. elegans germ line. J. Cell Biol. 192, 939–948 (2011).

12. K. E. Handwerger, J. A. Cordero, J. G. Gall, Cajal Bodies, Nucleoli, and Speckles in the Xenopus Oocyte Nucleus Have a Low-Density, Sponge-like Structure. Mol. Biol. Cell 16, 202–211 (2005).

13. L. Fu, et al., HIV-1 capsids enter the FG phase of nuclear pores like a transport receptor. Nature 626, 843–851 (2024).

14. V. Zila, et al., Cone-shaped HIV-1 capsids are transported through intact nuclear pores. Cell 184, 1032–1046.e18 (2021).

15. A. W. Folkmann, A. Putnam, C. F. Lee, G. Seydoux, Regulation of biomolecular condensates by interfacial protein clusters. Science 373, 1218–1224 (2021).

16. L.-H. Cai, S. Panyukov, M. Rubinstein, Mobility of nonsticky nanoparticles in polymer liquids. Macromolecules 44, 7853–7863 (2011).

17. S. Elbaum-Garfinkle, et al., The disordered P granule protein LAF-1 drives phase separation into droplets with tunable viscosity and dynamics. Proc. Natl. Acad. Sci. U. S. A. 112, 7189–7194 (2015).

18. B. S. Schuster, et al., Identifying sequence perturbations to an intrinsically disordered protein that determine its phase-separation behavior. Proc. Natl. Acad. Sci. 117, 11421–11431 (2020).

19. C. Iserman, et al., Genomic RNA Elements Drive Phase Separation of the SARS-CoV-2 Nucleocapsid. Mol. Cell 80, 1078–1091.e6 (2020).

20. C. A. Roden, et al., Double-stranded RNA drives SARS-CoV-2 nucleocapsid protein to undergo phase separation at specific temperatures. Nucleic Acids Res. 50, 8168–8192 (2022).

21. A. Savastano, A. Ibáñez de Opakua, M. Rankovic, M. Zweckstetter, Nucleocapsid protein of SARS-CoV-2 phase separates into RNA-rich polymerase-containing condensates. Nat. Commun. 11, 6041 (2020).

22. S. Rekhi, et al., Expanding the molecular language of protein liquid–liquid phase separation. Nat. Chem. 16, 1113–1124 (2024).

23. J. K. Armstrong, R. B. Wenby, H. J. Meiselman, T. C. Fisher, The Hydrodynamic Radii of Macromolecules and Their Effect on Red Blood Cell Aggregation. Biophys. J. 87, 4259– 4270 (2004).

24. B. S. Schuster, et al., Controllable protein phase separation and modular recruitment to form responsive membraneless organelles. Nat. Commun. 9, 2985 (2018).

25. M. Pizzinga, et al., Translation factor mRNA granules direct protein synthetic capacity to regions of polarized growth. J. Cell Biol. 218, 1564–1581 (2019).

26. R. Chouaib, et al., A Dual Protein-mRNA Localization Screen Reveals Compartmentalized Translation and Widespread Co-translational RNA Targeting. Dev. Cell 54, 773–791.e5 (2020).

27. D. M. Parker, L. P. Winkenbach, E. Osborne Nishimura, It’s Just a Phase: Exploring the Relationship Between mRNA, Biomolecular Condensates, and Translational Control. Front. Genet. 13 (2022).

28. Y. H. Tan, et al., A Nanoengineering Approach for Investigation and Regulation of Protein Immobilization. ACS Nano 2, 2374–2384 (2008).

29. I. Alshareedah, M. M. Moosa, M. Pham, D. A. Potoyan, P. R. Banerjee, Programmable viscoelasticity in protein-RNA condensates with disordered sticker-spacer polypeptides. Nat. Commun. 12, 6620 (2021).

30. L. M. Jawerth, et al., Salt-Dependent Rheology and Surface Tension of Protein Condensates Using Optical Traps. Phys. Rev. Lett. 121, 258101 (2018).

31. L. Jawerth, et al., Protein condensates as aging Maxwell fluids. Science 370, 1317–1323 (2020).

32. B. S. Schuster, D. B. Allan, J. C. Kays, J. Hanes, R. L. Leheny, Photoactivatable fluorescent probes reveal heterogeneous nanoparticle permeation through biological gels at multiple scales. J. Controlled Release 260, 124–133 (2017).

33. E. A. Nance, et al., A Dense Poly(Ethylene Glycol) Coating Improves Penetration of Large Polymeric Nanoparticles Within Brain Tissue. Sci. Transl. Med. 4, 149ra119–149ra119 (2012).

34. T. J. Böddeker, et al., Non-specific adhesive forces between filaments and membraneless organelles. Nat. Phys. 18, 571–578 (2022).

35. N. M. Green, “Avidin” in Advances in Protein Chemistry, C. B. Anfinsen, J. T. Edsall, F. M. Richards, Eds. (Academic Press, 1975), pp. 85–133.

36. F. Rico, A. Russek, L. González, H. Grubmüller, S. Scheuring, Heterogeneous and rate-dependent streptavidin–biotin unbinding revealed by high-speed force spectroscopy and atomistic simulations. Proc. Natl. Acad. Sci. 116, 6594–6601 (2019).

37. L.-H. Cai, S. Panyukov, M. Rubinstein, Hopping Diffusion of Nanoparticles in Polymer Matrices. Macromolecules 48, 847–862 (2015).

38. Y. Gu, M. E. Distler, H. F. Cheng, C. Huang, C. A. Mirkin, A General DNA-Gated Hydrogel Strategy for Selective Transport of Chemical and Biological Cargos. J. Am. Chem. Soc. 143, 17200–17208 (2021).

39. V. Grigorev, N. S. Wingreen, Y. Zhang, Conformational entropy of intrinsically disordered proteins bars intruders from biomolecular condensates. [Preprint] (2023). Available at: https://www.biorxiv.org/content/10.1101/2023.03.03.531005v1 [Accessed 20 July 2023].

40. Y. E. Guo, et al., Pol II phosphorylation regulates a switch between transcriptional and splicing condensates. Nature 572, 543–548 (2019).

41. D. L. J. Lafontaine, J. A. Riback, R. Bascetin, C. P. Brangwynne, The nucleolus as a multiphase liquid condensate. Nat. Rev. Mol. Cell Biol. 22, 165–182 (2021).

42. J. A. Riback, et al., Composition-dependent thermodynamics of intracellular phase separation. Nature 581, 209–214 (2020).

43. C. D. Reinkemeier, E. A. Lemke, Condensed, Microtubule-coating Thin Organelles for Orthogonal Translation in Mammalian Cells. J. Mol. Biol. 434, 167454 (2022).

44. L. L. J. Schoenmakers, et al., In Vitro Transcription–Translation in an Artificial Biomolecular Condensate. ACS Synth. Biol. 12, 2004–2014 (2023).

45. C. D. Reinkemeier, G. E. Girona, E. A. Lemke, Designer membraneless organelles enable codon reassignment of selected mRNAs in eukaryotes. Science 363, eaaw2644 (2019).

46. S. Nizzero, A. Ziemys, M. Ferrari, Transport Barriers and Oncophysics in Cancer Treatment. Trends Cancer 4, 277–280 (2018).

47. E. Blanco, H. Shen, M. Ferrari, Principles of nanoparticle design for overcoming biological barriers to drug delivery. Nat. Biotechnol. 33, 941–951 (2015).

48. M. J. Mitchell, et al., Engineering precision nanoparticles for drug delivery. Nat. Rev. Drug Discov. 20, 101–124 (2021).

49. A. Patel, et al., A Liquid-to-Solid Phase Transition of the ALS Protein FUS Accelerated by Disease Mutation. Cell 162, 1066–1077 (2015).

50. S. Mehta, J. Zhang, Liquid–liquid phase separation drives cellular function and dysfunction in cancer. Nat. Rev. Cancer 22, 239–252 (2022).

51. X. Tong, et al., Liquid–liquid phase separation in tumor biology. Signal Transduct. Target. Ther. 7, 1–22 (2022).

52. E. W. Martin, C. Iserman, B. Olety, D. M. Mitrea, I. A. Klein, Biomolecular Condensates as Novel Antiviral Targets. J. Mol. Biol. 436, 168380 (2024).

53. J. S. Oh, Y. Wang, D. J. Pine, G.-R. Yi, High-Density PEO-b-DNA Brushes on Polymer Particles for Colloidal Superstructures. Chem. Mater. 27, 8337–8344 (2015).

54. A. P. Thompson, et al., LAMMPS - a flexible simulation tool for particle-based materials modeling at the atomic, meso, and continuum scales. Comput. Phys. Commun. 271, 108171 (2022).

