## Supplementary Information for "Controlled and orthogonal partitioning of large particles into biomolecular condensates"

This PDF file includes:

Supporting text
Figures S1 to S13
Table S1
SI References

**Supporting Text**

*Sequences of proteins used in this paper*

Throughout this paper, RGG denotes residues 1-168 of LAF-1. N protein denotes SARS-CoV-2 nucleocapsid protein. $(GRGNSPYS)_{25}$ is an artificial intrinsically disordered protein containing 25 repeats of GRGNSPYS; for brevity, $(GRGNSPYS)_{25}$ is sometimes referred to as GRGNSPYS in the paper. SA denotes streptavidin and MBP denotes maltose-binding protein. All proteins include a hexahistidine tag at the N or C-terminus for immobilized metal affinity chromatography. Below, the main domains are color-coded; tags, linkers, and cut sites are not colored.

RGG
MESNQSNNGGSGNAALNRGGRYVPPHLRGGDGGAAAAASAGGDDRRGGAGGGGYRRGGG
NSGGGGGGGYDRGYNDNRDDRDNRGGSGGYGRDRNYEDRGYNGGGGGGGNRGYNNNRG
GGGGGYNRQDRGDGGSSNFSRGGYNNRDEGSDNRGSGRSYNNDRRDNGGDGLEHHHHHH

N protein
MHHHHHHENLYFQGMSDNGPQNQRNAPRITFGGPSDSTGSNQNGERSGARSKQRRPQGLPN
NTASWFTALTQHGKEDLKFPRGQGVPINTNSSPDDQIGYYRRATRRIRGGDGKMKDLSPRWYF
YYLGTGPEAGLPYGANKDGIIWVATEGALNTPKDHIGTRNPANNAAIVLQLPQGTTLPKGFYAEG
SRGGSQASSRSSSRSRNSSRNSTPGSSRGTSPARMAGNGGDAALALLLLDRLNQLESKMSGK
GQQQQGQTVTKKSAAEASKKPRQKRTATKAYNVTQAFGRRGPEQTQGNFGDQELIRQGTDYK
HWPQIAQFAPSASAFFGMSRIGMEVTPSGTWLTYTGAIKLDDKDPNFKDQVILLNKHIDAYKTFP
PTEPKKDKKKKADETQALPQRQKKQQTVTLLPAADLDDFSKQLQQSMSSADSTQA

$(GRGNSPYS)_{25}$
MRGSHHHHHHGSRSENLYFQGRSEFDPGRGNSPYSGRGNSPYSGRGNSPYSGRGNSPYSGR
GNSPYSGRGNSPYSGRGNSPYSGRGNSPYSGRGNSPYSGRGNSPYSGRGNSPYSGRGNSPY
SGRGNSPYSGRGNSPYSGRGNSPYSGRGNSPYSGRGNSPYSGRGNSPYSGRGNSPYSGRG
NSPYSGRGNSPYSGRGNSPYSGRGNSPYSGRGNSPYSGRGNSPYSGT

MBP-SA-RGG
MHHHHHHGGGSMKIEEGKLVIWINGDKGYNGLAEVGKKFEKDTGIKVTVEHPDKLEEKFPQVAA
TGDGPDIIFWAHDRFGGYAQSGLLAEITPDKAFQDKLYPFTWDAVRYNGKLIAYPIAVEALSLIYN
KDLLPNPPKTWEEIPALDKELKAKGKSALMFNLQEPYFTWPLIAADGGYAFKYENGKYDIKDVGV
DNAGAKAGLTFLVDLIKNKHMNADTDYSIAEAAFNKGETAMTINGPWAWSNIDTSKVNYGVTVLP
TFKGQPSKPFVGVLSAGINAASPNKELAKEFLENYLLTDEGLEAVNKDKPLGAVALKSYEEELVK
DPRIAATMENAQKGEIMPNIPQMSAFWYAVRTAVINAASGRQTVDEALKDAQTNSSSNNNNNNN
NNNLGENLYFQGMAEAGITGTWYNQLGSTFIVTAGADGALTGTYESAVGNAESRYVLTGRYDSA
PATDGSGTALGWTVAWKNNYRNAHSATTWSGQYVGGAEARINTQWLLTSGTTEANAWKSTLV
GHDTFTKVKPSAASLVPRGSMESNQSNNGGSGNAALNRGGRYVPPHLRGGDGGAAAAASAG
GDDRRGGAGGGGYRRGGGNSGGGGGGGYDRGYNDNRDDRDNRGGSGGYGRDRNYEDRG
YNGGGGGGGNRGYNNNRGGGGGYNRQDRGDGGSSNFSRGGYNNRDEGSDNRGSGRSYN
NDRRDNGGDGLEHHHHHH

*Derivation of LJ-based theoretical model*

To determine the change in energy from insertion of a particle into a condensed phase of protein, we consider two competing effects: 1) the displacement of favorable protein-protein interactions, and 2) the addition of favorable protein-bead interactions. We ignore bead-bead interactions because we are considering the case of insertion of a bead into a dense phase purely comprising protein. This assumption is likely one reason the theoretical boundary does not line up with the crossover point calculated from simulations.

To determine the energetic component of the transfer free energy, we define:

S1. $\Delta U_{\mathrm{transfer}} = \Delta U_{\mathrm{displace\ prot.}} + \Delta U_{\mathrm{prot-bead}}$

The displacement energy is assumed to be equal to the volume displaced by the bead, multiplied by the mean field internal energy of the pure protein liquid.

S2. $\Delta U_{\text{displace prot.}} = V_2 U_{(1)} \rho_{(1)}$

where $U_{(1)}$ is the per-particle energy of the system, and $\rho_{(1)}$ is the particle density. Note that these values are dependent on the conditions of the simulation, such as temperature. To obtain estimates for $U_{(1)}$ and $\rho_{(1)}$, we consider the pure dense phase of protein using the density profile from one case where protein-bead interactions are weak ($\varepsilon_{12} = 0.4$) and the concentration of beads inside the condensate is zero. This gives us a pure protein phase density of $\rho_{(1)} = 0.73$. The energy density of the dense phase of protein can be obtained by either running NVT simulation of pure protein in a box at density of 0.73, or can be interpolated from data already available through the NIST database (1).

The protein-bead interaction energy is assumed to be equal to a scaled-up version of a protein particle. Essentially, we take the per-particle energy from the pure protein case ($U_{(1)}$) and scale it up based on the increase in surface area ($A_2/A_1$), and the pairwise interaction energy ($\varepsilon_{12}/\varepsilon_1$). This gives:

S3. $\Delta U_{\text{prot-bead}} = U_{(1)} \left(\frac{A_2}{A_1}\right)\left(\frac{\epsilon_{12}}{\epsilon_1}\right)$

These are then combined to form equation 1 in the main text.

*Limitations of theory applied to simulations*

For small bead sizes, the partition coefficient is generally smaller than theory predicts. This may be due to several factors. First, the bead concentration at which the simulations were conducted is significantly higher than the theory accounts for, and thus the condensate environment is not purely protein. This effect would be particularly pronounced for cases with smaller beads, since the number of particles is greater than in cases with larger particles. Second, the theory assumes continuous energy density throughout the simulation box and that the attractive interactions between the bead and the protein particles will increase proportionally with respect to the surface area of the particle, which would be most valid at large bead sizes, but breaks down at small particle sizes. It also assumes that increased protein-bead interaction strength will not perturb the arrangement of protein molecules around the bead, which is also not true, but nontrivial to quantify. A more rigorous formulation may take into account the radial distribution function, which would need to be calculated separately for each case. Finally, in all cases, the concentration of protein particles in the low-density phase is significant, and will participate in attractive interactions with the beads, thus lowering the relative favorability of incorporating into the dense phase. This effect may be counteracted by reducing the temperature of the simulation, or by representing proteins as polymers. It may also be accounted for in the theoretical considerations by including a nonzero term for interactions in the dilute phase.

**Supplementary Figures**

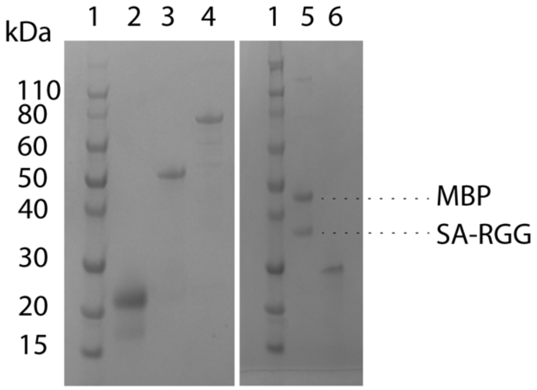


1. Ladder
2. RGG
3. N protein
4. MBP-SA-RGG, heated to 90°C
5. MBP-SA-RGG + TEV, heated to 90°C
6. $(GRGNSPYS)_{25}$

**Fig. S1. SDS-PAGE of purified proteins prepared for this study.** MBP-SA-RGG + TEV refers to MBP-SA-RGG that had been treated overnight by TEV protease to cleave off the MBP and liberate SA-RGG. MBP-SA-RGG and MBP-SA-RGG + TEV samples were heated at 90°C prior to running the gel to monomerize streptavidin (SA), which tetramerizes at room temperature. All other samples were prepared for running the gel by heating to 70°C.

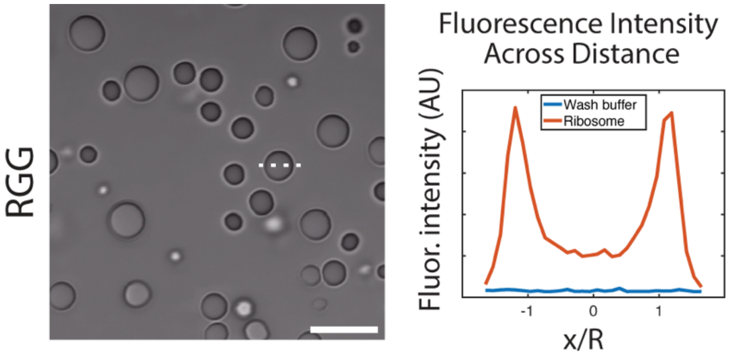


**Fig. S2. Partitioning of DL650-labeled ribosomes into RGG condensates, compared to partitioning of wash buffer following the ribosome labeling reaction.** (Left) Partitioning of wash buffer into RGG. The partitioning of DL650-labled ribosomes into RGG is shown in Fig. 1B. (Right) Fluorescence intensity (not normalized) across distance, comparing DL650-ribosomes (red) and wash buffer (blue). The purpose of this comparison is to show that unbound dye was thoroughly removed from the DL650-labeled ribosomes, so the fluorescent signal shown in Fig. 1B is from the labeled ribosomes and not unbound dye.

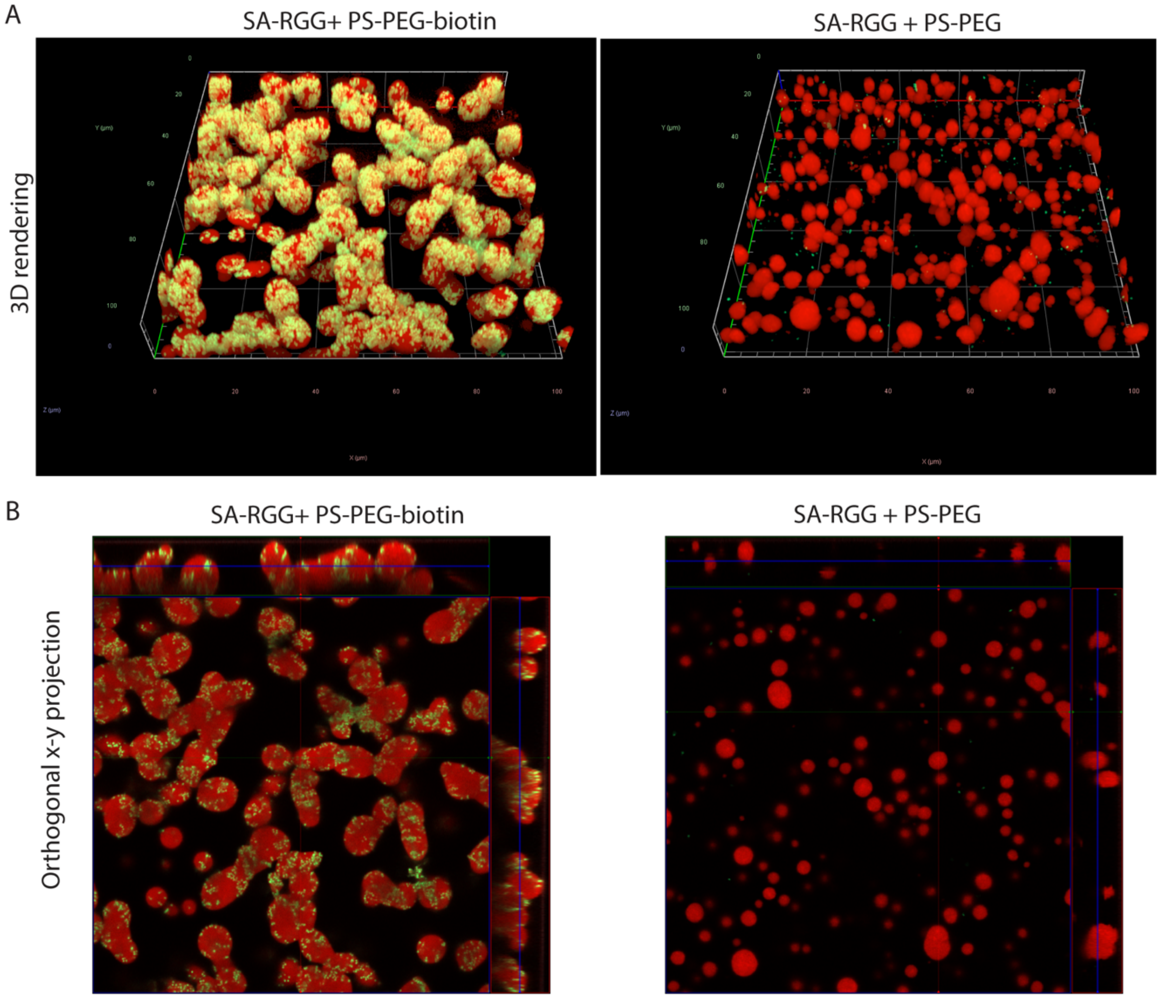


**Fig. S3. Confocal microscopy of SA-RGG + PS-PEG-biotin vs. SA-RGG + PS-PEG.** Rhodamine (red) was added to visualize the SA-RGG condensates. Nanoparticles (500 nm diameter) are green. (A) 3D renderings from z-stacks of SA-RGG + PS-PEG-biotin (left) and SA-RGG + PS-PEG (right). (B) Orthogonal projections showing xy, xz, and yz planes.

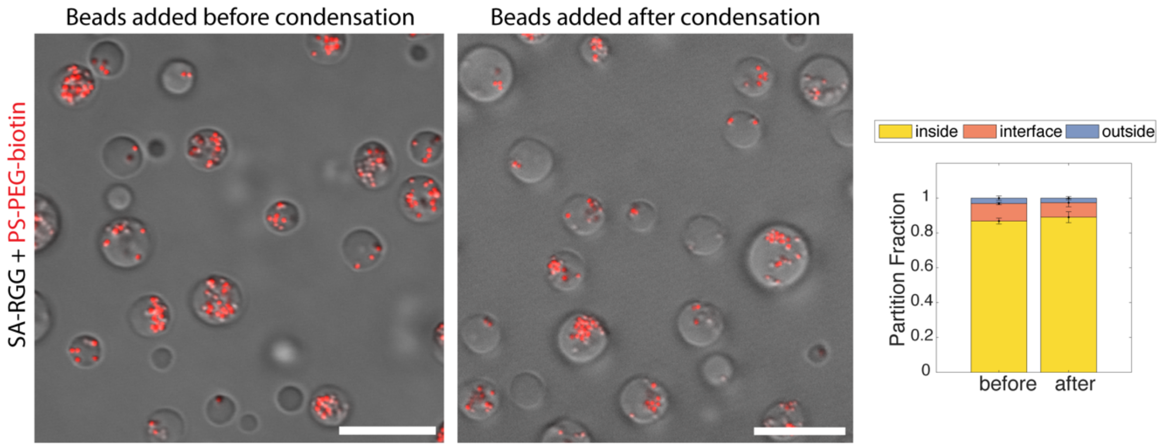


**Fig. S4. PS-PEG-biotin beads (500 nm) added to SA-RGG before vs. after phase separation of SA-RGG.** The partition fraction of the beads into SA-RGG condensates is similar whether the beads were added before or after condensate formation.

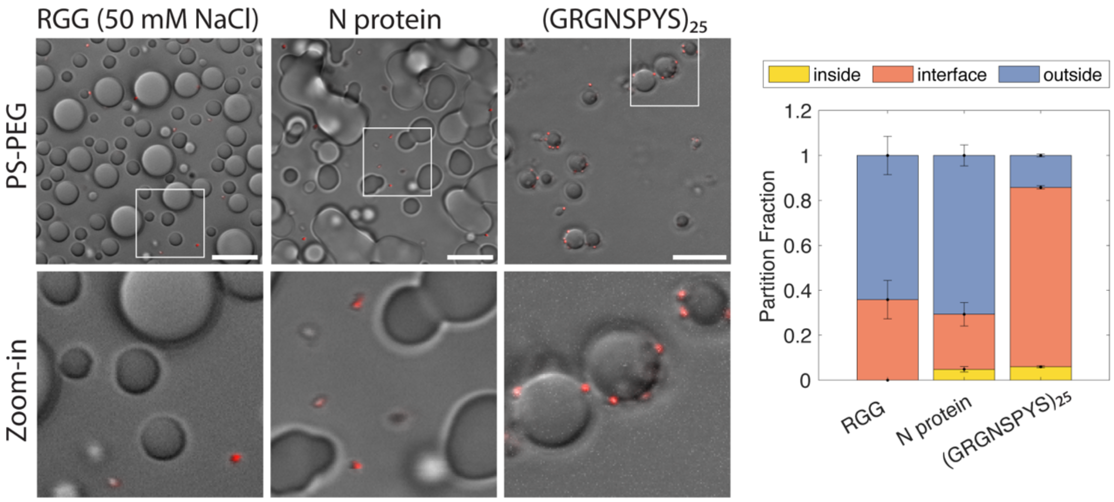


**Fig. S5. PS-PEG beads (500 nm) are excluded from RGG, N protein, and $(GRGNSPYS)_{25}$ condensates.** Here, RGG was prepared in 50 mM NaCl buffer, and the other two proteins were prepared in 150 mM NaCl buffer. (Left) Microscopy images. (Right) Partition fractions were quantified. Error bars represent SEM with n = 10.

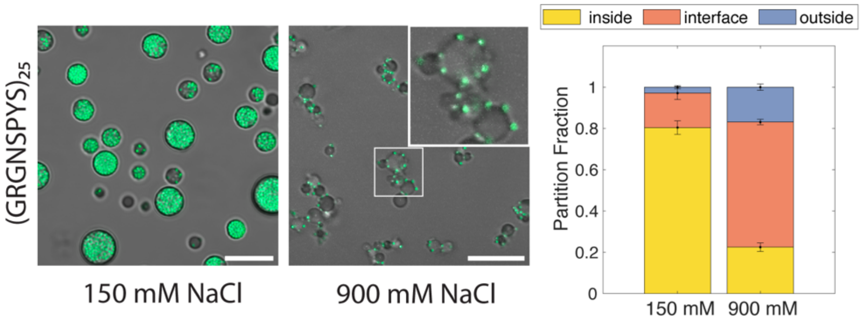


**Fig. S6. GRGNSPYS condensates with 500 nm PS-PEG-polyA20 beads (green) in 150 mM NaCl vs. 900 mM NaCl buffers** (both with 20 mM Tris, pH 7.5)**.** (Left) Micrographs show beads are excluded from condensates at high salt concentration. Inset in 900 mM NaCl image shows zoom-in of boxed region. (Right) Partition fractions were quantified. Error bars represent SEM with n = 10.

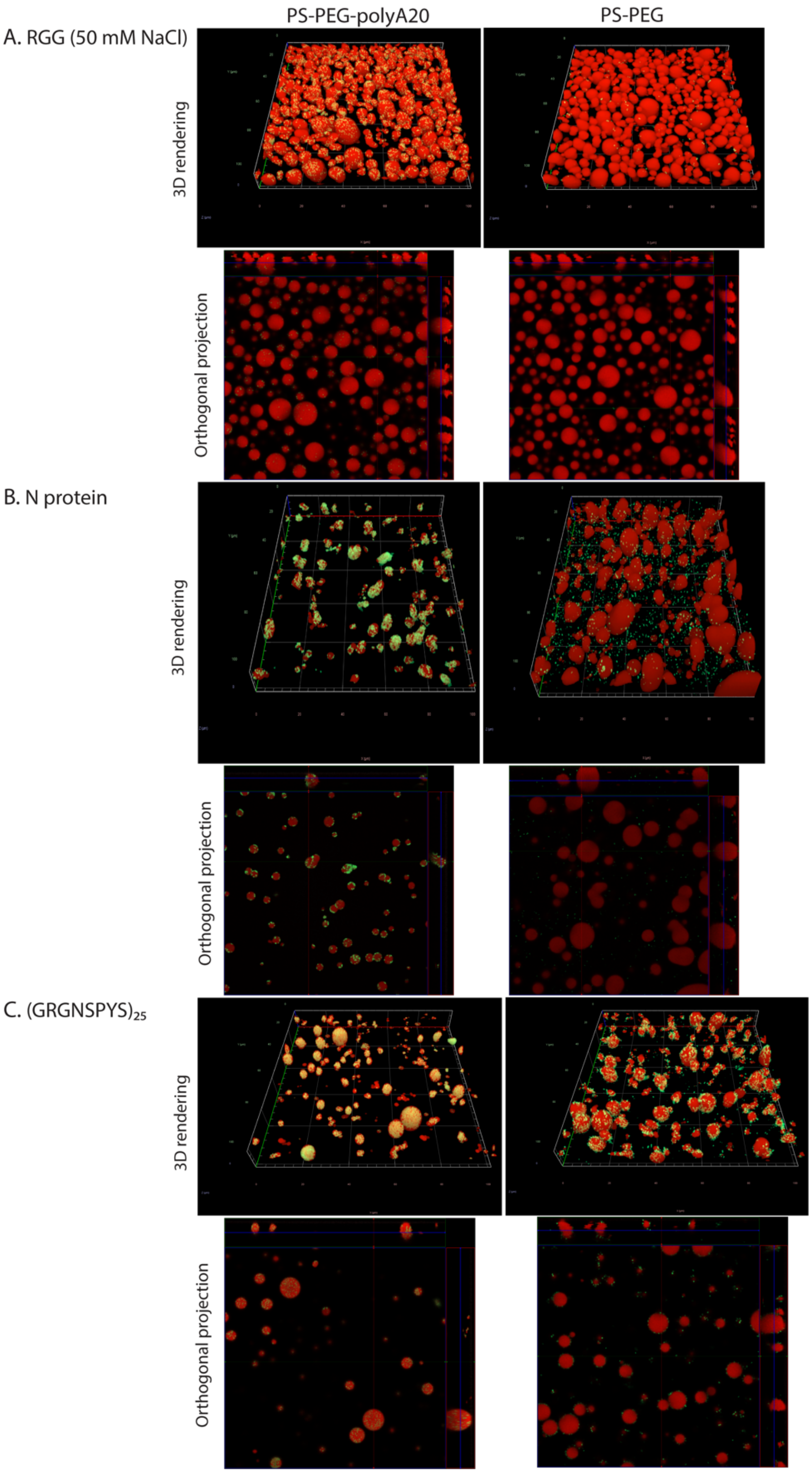


**Figure S7. Confocal microscopy of condensates with PS-PEG-polyA20 vs. PS-PEG beads.** Rhodamine (red) was added to visualize the condensates. Nanoparticles (500 nm diameter) are green. PS-PEG in this figure refers to PS-PEG-azide, where the free end of the PEG has an azide functional group, which was subsequently used for click chemistry reaction with DBCO-polyA20**.** (A) LAF-1 RGG at 50 mM NaCl. (B) N protein at 150 mM NaCl. (C) GRGNSPYS at 150 mM NaCl. For each protein, 3D renderings and orthogonal projections are shown.

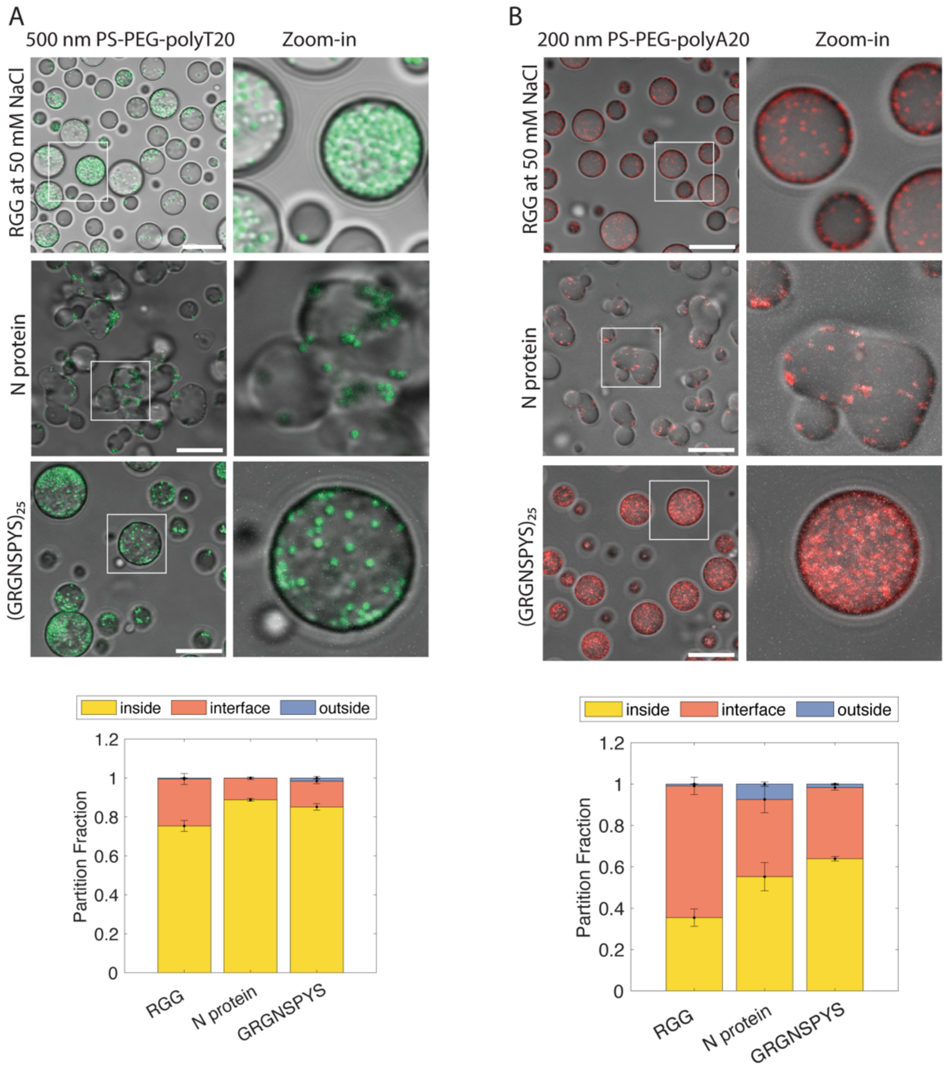


**Figure S8. Additional oligonucleotide sequence (polyT20) and bead size (200 nm) tested for PS-PEG-oligo partitioning into condensates.** (A) 500 nm PS-PEG-polyT20 beads partition into RGG (at 50 mM NaCl), N protein (at 150 mM NaCl), and GRGNSPYS (at 150 mM NaCl). (B) 200 nm PS-PEG-polyA20 beads also partition. Error bars represent SEM with n ≥ 5.

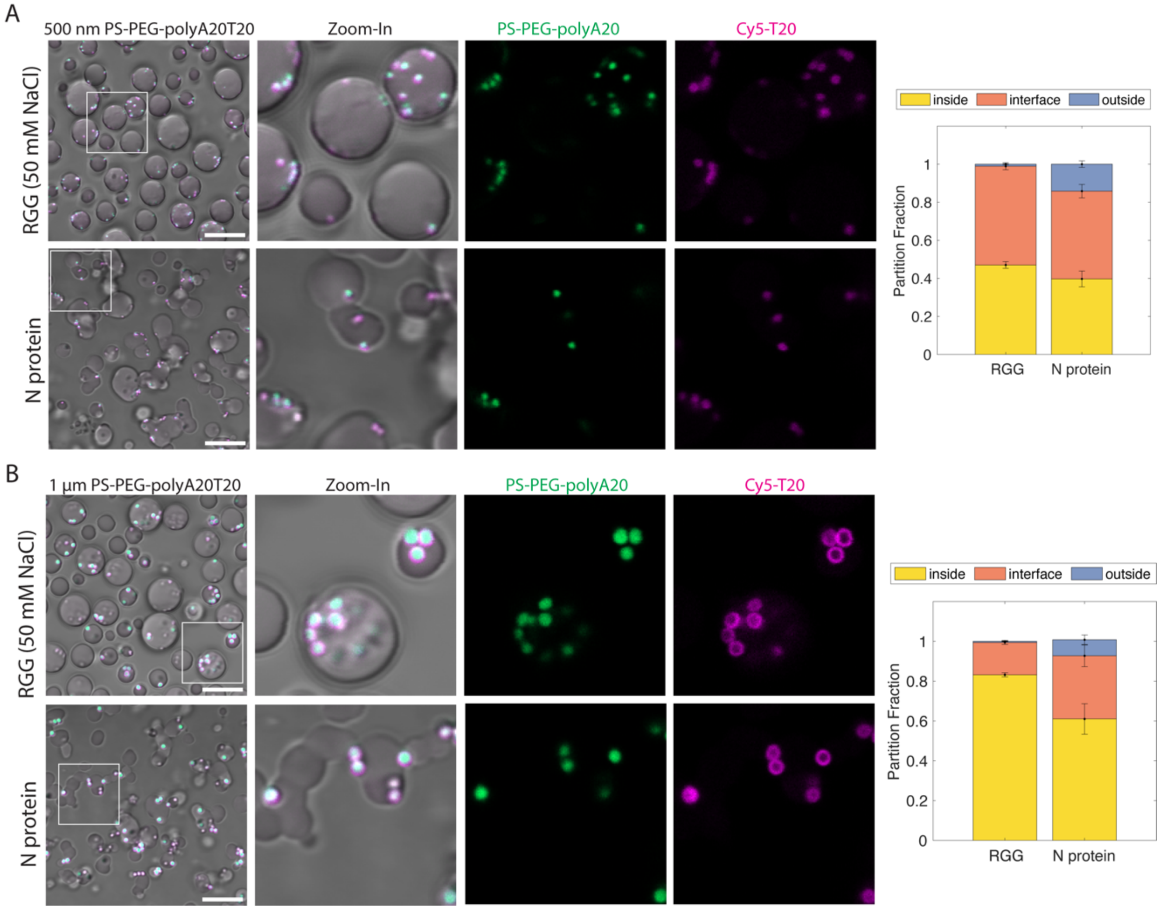


**Fig. S9. Beads conjugated with double-stranded oligonucleotides partition into condensates.** RGG is in 50 mM NaCl buffer and N protein in 150 mM NaCl buffer. polyA20T20 beads had been prepared as follows: polyA20 was conjugated to PS-PEG beads (green) by click chemistry, then Cy5-polyT20 was annealed to the PS-PEG-polyA20 beads. Colocalization of the Cy5 signal with the beads demonstrates successful annealing between polyT20 and PS-PEG-polyA20 to form PS-PEG-polyA20T20. (A) Partitioning of 500 nm PS-PEG-polyA20T20. (B) Partitioning of 1 µm PS-PEG-polyA20T20. Scale bars represent 10 µm. Error bars represent SEM with n = 10.

PS-PEG-75%A20

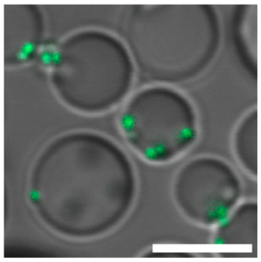

**Fig. S10. PS-PEG-75%polyA20 beads (500 nm) adhere to the interface of RGG condensates** (in 50 mM NaCl buffer)**.** Partitioning is quantified in Fig. 4E.

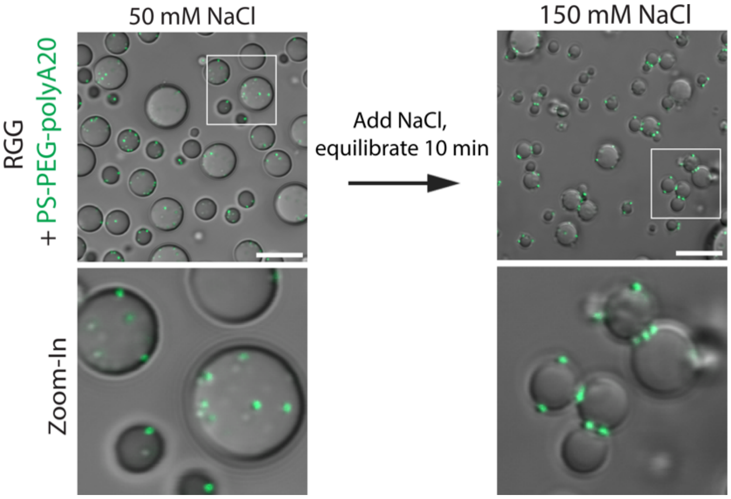


**Figure S11. Reversing partitioning by increasing salt concentration.** PS-PEG-polyA20 particles (green) partition into RGG condensates at 50 mM NaCl. When NaCl concentration is raised to 150 mM, previously partitioned beads become excluded from condensates. Scale bars, 10 μm. (Right) Partition fractions quantified. Error bars represent SEM with $n \geq 9$.

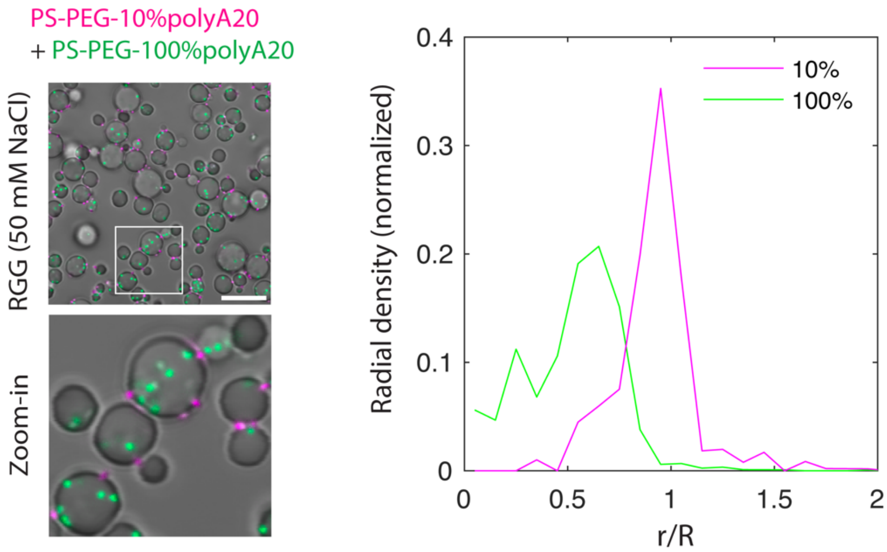


**Figure S12. Selective targeting of condensate interior vs. interface based on oligo density on bead surface.** Condensates are RGG at 50 mM NaCl**.** Bead diameter is 500 nm. Micrographs show PS-PEG-10%polyA20 beads (magenta) adsorb to the condensate interface and PS-PEG-100%polyA20 beads (green) are enriched inside condensates. (Right) Radial density profile shows bead concentration as a function of normalized distance from condensate center. (Scale bar, 10 $\mu$m.)

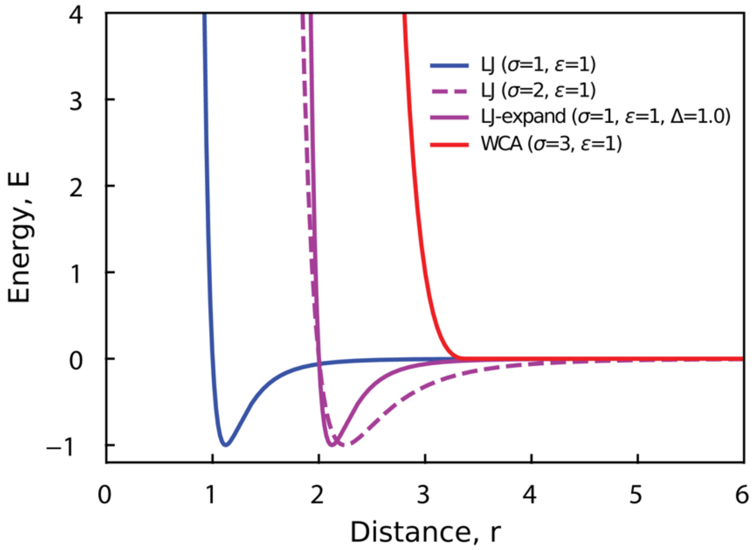


**Figure S13. Energy functions for pairwise interactions between spherical particles used in molecular dynamics simulations.** As an example, we are showing functional forms for the case where bead size is 3 (protein size is always 1, so for protein-bead interactions in this example, the average $\sigma = 2$ is used). The protein-protein interactions are a simple LJ functional form, while protein-bead interactions are handled by an LJ-expand functional form, which keeps the shape of the energy well the same as with protein-protein interactions, but shifts it outward to increase the effective size of the beads. For comparison, the LJ functional form for the same size is shown as a dashed line, illustrating the widening of the energy well. Finally, bead-bead interactions were treated repulsively using a Weeks-Chandler-Andersen (WCA) potential.

| Particle | Size (nm) |
|---|---|
| 200 nm PS | 223 ± 3.71 |
| 200 nm PS-PEG-azide | 199 ± 0.06 |
| 500 nm PS | 567 ± 24.6 |
| 500 nm PS-PEG-azide | 619 ± 12.3 |
| 1 μm PS | 1190 ± 142 |
| 1 μm PS-PEG-azide | 1050 ± 51.9 |
| 500 nm PS-PEG-polyA20 | 646 ± 42.9 |
| 500 nm PS-PEG-polyA40 | 541 ± 6.01 |
| 500 nm PS-PEG-10% polyA20 | 598 ± 13.1 |
| 500 nm PS-PEG-75% polyA20 | 622 ± 9.07 |
| 500 nm PS-PEG-polyA20T20 | 642 ± 5.64 |
| 1 μm PS-PEG-polyA20T20 | 1190 ± 146 |
| 100 nm PS-PEG-30% biotin | 140 ± 0.56 |
| 200 nm PS-PEG-20% biotin | 211 ± 0.09 |
| 500 nm PS-PEG-10% biotin | 578 ± 13.4 |
| 500 nm PS-PEG-100% biotin | 647 ± 30.8 |
| 200 nm PS-PEG-polyA20 | 226 ± 2.54 |

| Particle | Charge (mV) |
|---|---|
| 200 nm PS | -51.2 ± 5.34 |
| 200 nm PS-PEG-azide | 0.11 ± 0.4 |
| 500 nm PS | -65.9 ± 1.68 |
| 500 nm PS-PEG-azide | -0.51 ± 0.66 |
| 500 nm PS-PEG-10% polyA20 | -4.7 ± 5.81 |
| 500 nm PS-PEG-75% polyA20 | -29.2 ± 1.7 |
| 500 nm PS-PEG-100% polyA20 | -34 ± 4.67 |
| 500 nm PS-PEG-polyA5 | -15.75 ± 5.3 |
| 500 nm PS-PEG-polyA40 | -36.6 ± 5.8 |
| 500 nm PS-PEG-polyA20T20 | -28.4 ± 0.969 |

**Table S1.** Size and zeta potential measurements of beads used in this study. Particle size was measured by dynamic light scattering. Zeta potential was measured by laser doppler electrophoresis.

**Supplementary Methods**

*Cloning*

RGG, N protein, and MBP-SA-RGG were cloned into a pET vector in-frame with 6xHis-tag using NEBuilder HiFi DNA Assembly (New England BioLabs) and appropriate primers and PCR products or synthetic gene fragments (gBlocks; IDT). Gene sequences was verified using whole plasmid, long-read sequencing (Plasmid-EZ, GENEWIZ). $(GRGNSPYS)_{25}$ was cloned by Genscript into a pQE80L vector.

*Protein expression and purification*

RGG, N protein, and MBP-SA-RGG were expressed and purified using methods previously described (2, 3). In brief: Plasmids were transformed into BL21 (DE3) competent *E. coli* cells and recombinantly expressed and purified by immobilized metal affinity chromatography. Proteins were washed with 500 mM NaCl, 20 mM Tris-HCl, 20 mM imidazole, pH 7.5 buffer and eluted with 500 mM NaCl, 20 mM Tris-HCl, 500 mM imidazole, pH 7.5 buffer. N protein was purified in a similar method, but proteins bound to the column were washed with 3 M NaCl, 20 mM Tris-HCl, 20 mM imidazole buffer. Only eluted N protein fractions with A260/A280 ratio < 0.7 were used, to avoid DNA and RNA contamination.

GRGNSPYS was transformed into *E. coli* M15-[pREP4] strain and recombinantly expressed and purified using previously described methods (2).

Purified proteins were dialyzed overnight using 7 kDa MWCO membranes (Slide-A-Lyzer G2, ThermoFisher) in appropriate buffers. RGG was dialyzed in 150 mM NaCl, 20 mM Tris, pH 7.5, at 45 ℃ to inhibit phase separation. MBP-SA-RGG was dialyzed at room temperature and sterile filtered through a 0.45 μm filter (SLHPX13NL; MilliporeSigma). N protein aliquots with A260/A280 ratio < 0.7 were dialyzed into 300 mM NaCl, 20 mM Tris buffer, pH 7.5 at room temperature. Dialyzed RGG and N protein aliquots were flash frozen and stored at -80 °C. Dialyzed MBP-SA-RGG aliquots were stored at 4 °C.

SDS-PAGE was carried out on all proteins to confirm purity using NuPAGE 4-12% Bis-Tris gels (Invitrogen) followed by incubation with Coomassie stain (GelCode™ Blue Safe Protein Stain; ThermoFisher). Protein concentrations were measured using NanoDrop One Microvolume Spectrophotometer (ThermoFisher). RGG and GRGNSPYS concentration were measured in final 4 M urea buffer to prevent phase separation.

*Preparation and storage of purchased proteins and small molecules*

Lyophilized polyA RNA (P9403; Sigma) used to induce N protein phase separation was resuspended in MilliQ water to 10 mg/mL, flash frozen, and stored at -80 °C. Antibodies were purchased from ThermoFisher: SARS-CoV-2 Nucleocapsid protein rabbit polyclonal IgG (catalog #: SARS-COV2-N-FITC) and Rabbit IgG Isotype Control, FITC (catalog # 11-4614-80) and stored according to the manufacturer's recommendations. Ribosomes were purchased from New England Biolabs (*E. coli* Ribosome) and fluorescently labeled with DyLight 650 NHS Ester (DL650) (catalog # 62265; ThermoFisher) using a previously described protocol (3). After the labeling reaction, ribosomes were thoroughly washed in a centrifugal ultrafiltration device (UFC501096; Sigma) until no fluorescence was observed from condensates mixed with the wash buffer flow-through. Aliquots of labeled ribosomes were flash frozen and stored at -80 °C. Biotin-4-fluorescein was purchased from AAT Bioquest, resuspended in DMSO at 10 mg/mL, and stored at -20 °C. Dextrans of various sizes were purchased from Sigma (R8881, 42874, R9379), resuspended in MilliQ water at 50 mg/mL, and stored at 4 °C.

*Oligonucleotide preparation*

Oligonucleotide strands were synthesized using a K&A H-2 synthesizer on Glen UnySupport™ 1000 (Glen Research) with TWIST columns (Glen Research) on a 10 µmol scale. All phosphoramidites and oligonucleotide synthesis reagents were purchased from Glen Research and used as received. The synthesis utilized dA-CE phosphoramidite, dT-CE phosphoramidite, Ac-dC-CE phosphoramidite, and dmf-dG-CE phosphoramidite to incorporate standard ATCG bases. 5' Cy5 and DBCO modifications were done using Cyanine 5 phosphoramidite and 5'-DBCO-TEG phosphoramidite, respectively. Synthesis was performed using 0.25 M 5-ethylthio-1H-tetrazole (ETT) in anhydrous acetonitrile as the activator and 3% TCA/DCM as the deblocking reagent. The capping step was carried out by mixing tetrahydrofuran/pyridine/acetic anhydride (Cap Mix A) and 16% 1-methylimidazole in tetrahydrofuran (Cap Mix B). Oxidation was carried out using 0.02 M iodine in tetrahydrofuran/water/pyridine for all non-DBCO containing strands (with a 40-second oxidation time) and 0.5 M CSO in anhydrous acetonitrile for DBCO-containing strands (with a 3-minute oxidation time). Coupling times were set at 55 seconds for standard ATCG bases, 3 minutes for Cyanine 5 phosphoramidite, and 10 minutes for 5'-DBCO-TEG phosphoramidite. Strands were deprotected and cleaved using a 35% ammonia solution (VWR) for 17 hours at 55 °C for non-Cy5-containing strands, or 17 hours at room temperature for Cy5-containing strands. All oligonucleotides were purified by reverse-phase HPLC (SHIMADZU LC-20AR) using a ZORBAX StableBond 300 C18, 250 X 9.4 mm I.D., 5 μm as the stationary phase, with a flow rate of 3 mL/min, and acetonitrile and 0.06 M triethylamine acetate (TEAA) aqueous buffer as the mobile phase. Once purified, all oligonucleotides were lyophilized and characterized using ESI-HRMS (Waters SYNAPT G2-Si Mass Spectrometry). Purity was confirmed via analytical HPLC using a C18 column (150 X 4.6 mm I.D.) from YMC CO., LTD.

A complete list of the synthesized oligonucleotides is shown below.

| | DNA sequence |
|---|---|
| Cy5-polyA20 | 5′ Cy5-A20 3′ |
| Cy5-polyT20 | 5′ Cy5-T20 3′ |
| polyA5-DBCO | 5′ DBCO-A5 3′ |
| polyA20-DBCO | 5′ DBCO-A20 3′ |
| polyA40-DBCO | 5′ DBCO-A40 3′ |
| polyT20-DBCO | 5′ DBCO-T20 3′ |

*Bead preparation*

Particle PEGylation reactions were performed using previously described methods optimized by Nance et al (4). Amine-terminated PEG polymers were conjugated to carboxyl-modified polystyrene beads using NHS-EDC chemistry in pH 8.2, 200 mM borate buffer (product numbers in table below). All bead modifications were done in 1.5 mL tubes (1615-5500; USA Scientific) prepared by coating with 5% Pluronic F-127 for a minimum of 1 hour, followed by rinsing once with 1 mL MilliQ water, to minimize bead loss. Depending on the beads' COOH density (as measured by the manufacturer), the mass of PEG, N-hydroxysulfosuccinimide (sulfo-NHS), 1-ethyl-3-(3-dimethylaminopropyl)carbodiimide hydrochloride (EDC), and the borate buffer volume were determined below. Reactions were incubated for a minimum of 4 hours at room temperature on a rotary shaker.

To prepare PS-PEG-biotin beads, biotin-PEG-amine was used. Density of biotin around beads were controlled by mixing biotin-PEG-amine with mPEG-amine in appropriate ratios during PEGylation. For example, to prepare PS-PEG-15% biotin, 15 weight % biotin-PEG-amine was mixed with 85 weight % mPEG-amine. To compare biotin density across different bead sizes, first the bead concentration was quantified using a plate reader (Molecular Devices, spectraMax M2) in a 96-well black bottom plate (Fisher). Pierce™ Fluorescence Biotin Quantitation Kit (ThermoFisher) was used to determine the biotin density on PS-PEG-biotin beads and measured using the same plate reader. The biotin standard curve was generated using the kit with a 488 nm excitation. Bead concentration standard curves were generated for each bead size with unmodified red beads in the plate reader using a 561 nm excitation. The bead concentrations of biotinylated red beads were measured and analyzed using the same excitation. Then, the biotinylated beads were analyzed for biotin density using 488 nm excitation. The biotin density for each bead sample was calculated using the 488 nm and 561 nm standard curves.

To prepare PS-PEG-oligonucleotide beads, the beads were first PEGylated using azide-PEG-amine. Then, oligonucleotide conjugation reaction was performed through strain-promoted azide-alkyne cycloaddition (SPAAC) click chemistry (5). DBCO-functionalized oligonucleotides were added to PS-PEG-azide in a 1:1 ratio with respect to the COOH groups in 150 mM NaCl, pH 7.5 and incubated on a rotary shaker for 24 hours. Density of oligonucleotide on the beads was controlled beginning in the PEGylation step by mixing azide-PEG-amine with mPEG-amine in appropriate ratios. For example, to prepare PS-PEG-75% polyA20, 75 weight % azide-PEG-amine was mixed with 25 weight % mPEG-amine during the PEGylation reaction.

Beads were washed twice with MilliQ water by centrifugation, according to the tabulated centrifugation speeds and durations by size below. Beads were then resuspended in MilliQ water and stored at 4 °C. Beads were vortexed briefly prior to mixing with proteins for microscopy to prevent bead aggregation.

All PEG polymers were purchased from Creative PEG Works in 5k MW.

| Bead | PEG polymer | Polymer Catalog Number |
|---|---|---|
| PS-PEG-biotin | biotin-PEG-amine | PJK-1942 |
| PS-PEG-oligonucleotide | azide-PEG-amine | PHB-1884 |
| PS-PEG | mPEG-amine | PLS-268 |

PEGylation reaction conditions for Fluospheres (ThermoFisher) carboxylate-modified polystyrene microspheres are shown below.

| | Red Fluorescent | | | | Yellow-Green Fluorescent | | | |
|---|---|---|---|---|---|---|---|---|
| Size | 100 nm | 200 nm | 500 nm | 1 μm | 100 nm | 200 nm | 500 nm | 1 μm |
| Product number | F8887 | F8887 | F8887 | F8887 | F8888 | F8888 | F8813 | F8888 |
| Volume of beads | 50 μL | 50 μL | 50 μL | 50 μL | 50 μL | 50 μL | 50 μL | 50 μL |
| Mass of PEG (mg) | 6.18 | 5.34 | 0.31 | 0.62 | 3.17 | 5.25 | 2.69 | 1.98 |
| Mass of NHS (mg) | 2.68 | 2.32 | 0.13 | 0.27 | 1.37 | 2.28 | 1.17 | 0.86 |
| Vol. borate buffer (μL) | 800 | 800 | 400 | 400 | 800 | 800 | 800 | 800 |
| Mass of EDC (mg) | 11.5 | 14.38 | 7.19 | 7.19 | 5.75 | 14.38 | 14.38 | 28.76 |
| Centrifugation speed (x g) | 21,000 | 18,000 | 15,000 | 10,000 | 21,000 | 18,000 | 15,000 | 10,000 |
| Centrifugation time | 25 min | 15 min | 12 min | 10 min | 25 min | 15 min | 12 min | 10 min |

PEGylation reaction conditions for Fluorescent Carboxyl Polystyrene beads from Bangs Labs are shown below.

| | Green Fluor. | Far Red Fluor. |
|---|---|---|
| Size | 500 nm | 500 nm |
| Product number | FCDG005 | FCFR005 |
| Volume of beads | 50 μL | 50 μL |
| Mass of PEG (mg) | 2.44 | 2.44 |
| Mass of NHS (mg) | 1.06 | 1.06 |
| Volume of borate buffer (μL) | 800 | 800 |
| Mass of EDC (mg) | 14.38 | 14.38 |
| Centrifugation speed (x g) | 15,000 | 15,000 |
| Centrifugation time | 12 min | 12 min |

*Microscopy*

Imaging was carried out on a Zeiss LSM900 confocal microscope with an Axio Observer 7 inverted stand and using a 63x plan-apochromatic, oil-immersion objective with 1.4 numerical aperture (NA). Transmitted light images were collected with an ESID module (0.55 NA condenser). Green beads, biotin-4-fluorescein, and fluorescein-labeled antibodies were excited with a 488 nm laser; red beads and rhodamine labeled dextran with a 561 nm laser; and DL650 labeled ribosomes, AF650 labeled N protein, far-red fluorescent beads, and Cy5-labeled oligonucleotides with a 640 nm laser.

RGG samples were imaged at 1 mg/mL in 150 mM NaCl, 20 mM Tris, pH 7.5, unless otherwise stated. Protein was thawed at 45 °C immediately before imaging to prevent aggregation. N protein was prepared at 45 µM in 150 mM NaCl, 20 mM Tris, pH 7.5 buffer and mixed with 0.5 mg/mL polyA RNA (P9403; Sigma-Aldrich) immediately prior to imaging to induce phase separation. GRGNSPYS was imaged at 18 µM in 1x PBS buffer.

MBP-SA-RGG at 4 mg/mL was reacted with 0.02 mg/mL final TEV protease concentration for 30 minutes to induce phase separation by removing the MBP tag. This was done in Pluronic F-127 coated microscope dishes. Typically, phase separation was induced before beads were added. For control samples where beads were added before phase separation, beads were added to MBP-SA-RGG before TEV protease treatment.

Beads were added at 0.01 volume % unless otherwise specified. For antibody partitioning experiments, antibodies were mixed with protein to final 0.33 μM. For dextran partitioning experiments, dextrans were mixed with proteins at final 0.5 g/L. For oligonucleotide partitioning experiments, Cy5-polyA20 was mixed with proteins at final 5.5 μM. For biotin-4-fluorescein partitioning experiments, biotin-4-fluorescein was mixed with SA-RGG at final 0.1 g/L.

Samples were plated on 16-well glass-bottom dishes (#1.5 glass thickness; Grace Bio-Labs) after pre-treating the glass with a solution of 5% Pluronic F-127 (Sigma-Aldrich) for a minimum of 10 minutes and then rinsing with DI water. Plated samples were equilibrated for 10 minutes before imaging.

*Image analysis*

Image analysis and quantification were performed using custom codes in MATLAB R2023b. Condensates were identified in each image by segmentation or by the circular Hough Transform. Bead positions were identified as centroids of bright spots in the appropriate fluorescence channel. The position of fluorescent particles with respect to the condensates were classified as inside, interface, or outside the condensates. For dextrans and antibodies, the partition coefficient was measured as the average fluorescence intensity inside the condensates, divided by the average fluorescence intensity outside the condensates (after subtracting background fluorescence measured from a blank sample). Bar graphs, line scans, and radial concentration profiles were also generated in MATLAB.

*Particle size and zeta potential measurements*

Bead size and zeta potential were measured using a Malvern Zetasizer Ultra. Bead size was measured using 1 μL of bead sample diluted into 1 mL MilliQ water in a polystyrene cuvette. Bead zeta potential values were measured using 1 μL of bead sample in 700 μL of 0.1x PBS buffer in a folded capillary zeta cell (DTS1070; Malvern).

*Molecular dynamics simulations*

Simulations were carried out on systems of Lennard-Jones (LJ) particles using the LAMMPS simulation engine (6). In each system, we simulate two types of particles, 1) protein, and 2) an attractive client molecule, simply referred to as a "bead." Protein-protein interactions were handled using a simple LJ functional form:

1) $$\mathrm{E(r)} = 4\epsilon_1 \left[\left(\frac{\sigma_1}{\mathrm{r}}\right)^{12} - \left(\frac{\sigma_1}{\mathrm{r}}\right)^{6}\right],\ \mathrm{r} < \mathrm{r_{cut}}$$

where $\epsilon_1 = 1$ is the depth of the energy well, or the strength of attractive interactions between two protein particles, and $\sigma_1 = 1$ is the characteristic distance of the interactions, or the size of the LJ particle representing protein molecules.

Bead-bead interactions were treated as softly repulsive, and handled using a Weeks-Chandler-Anderson functional form:

2) $$\mathrm{E(r)} = \begin{cases} 4\epsilon_2 \left[\left(\frac{\sigma_2}{\mathrm{r}}\right)^{12} - \left(\frac{\sigma_2}{\mathrm{r}}\right)^{6}\right] + \epsilon_2\,, & \mathrm{r} < 2^{\frac{1}{6}}\sigma_2 \\ 0 & ,\ \mathrm{r} \geq 2^{\frac{1}{6}}\sigma_2 \end{cases}$$

where $\epsilon_2 = 1$ is the scaling of repulsive interactions between beads and $\sigma_2$ is the size of a bead, which is treated as a free variable to test the size-dependence of partitioning of beads into a condensate of protein particles.

Since the surface chemistry of beads and their interactions with protein should be similar at different sizes, for protein-bead interactions we utilize the lj-expand force field, which keeps the width of the energy well identical between all bead sizes, and simply shifts it outward further from the center of the bead to increase the bead's size:

3) $E(r) = 4\epsilon_{12}\left[\left(\frac{\sigma_1}{r-\Delta_{12}}\right)^{12} - \left(\frac{\sigma_1}{r-\Delta_{12}}\right)^{6}\right],\ r < r_{cut} + \Delta_{12}$

where $\Delta_{12} = \frac{\sigma_2+\sigma_1}{2} - 1$ is the offset value used to shift the LJ functional form outward from the center of the bead and $\epsilon_{12}$ is the interaction energy between protein and beads. For all systems tested, protein particles were assigned default LJ parameters: $\sigma_1 = 1$, $\epsilon_1 = 1$, and mass = 1, while beads were assigned a range of $\epsilon_{12}$ and $\sigma_2$ parameters. Masses of beads were increased to keep density constant at 1, and scaling of repulsive interactions was set to be similar to that of the protein particles by setting $\epsilon_2 = 1$. Functional forms used for each interaction are shown in Fig. S13.

Simulations were conducted using 51200 protein particles, and enough beads to reproduce a 5:1 protein:bead mass ratio. We conducted simulations in slab geometry with a box of size 20x20x200 $\sigma^3$ for 10 million MD steps using Langevin dynamics at constant temperature: $T^* = 0.9$. The condensed phase was centered in the box using the methodology of Jung et al. (7) and density profiles were calculated by binning the elongated z-dimension into 500 bins and calculating the density of particles at each frame and discarding the first 2 million steps as equilibration of the system. Partition coefficients were calculated by first calculating equilibrium concentrations of beads inside and outside the slab. The boundaries of the condensate and external regions were determined by selecting a sub-region of the system where bead concentration is relatively level, and averaging values across all bins within this range. Importantly, some systems had significant interfacial enrichment, and thus the size of the box used for this calculation was smaller in those cases.

**SI References**


1. Daniel Siderius, NIST Standard Reference Simulation Website - SRD 173. National Institute of Standards and Technology. https://doi.org/10.18434/MDS2-232. Deposited 1 September 2017.

2. S. Rekhi, *et al.*, Expanding the molecular language of protein liquid–liquid phase separation. *Nat. Chem.* **16**, 1113–1124 (2024).

3. L. L. J. Schoenmakers, *et al.*, In Vitro Transcription–Translation in an Artificial Biomolecular Condensate. *ACS Synth. Biol.* **12**, 2004–2014 (2023).

4. E. A. Nance, *et al.*, A Dense Poly(Ethylene Glycol) Coating Improves Penetration of Large Polymeric Nanoparticles Within Brain Tissue. *Sci. Transl. Med.* **4**, 149ra119-149ra119 (2012).

5. J. S. Oh, Y. Wang, D. J. Pine, G.-R. Yi, High-Density PEO-b-DNA Brushes on Polymer Particles for Colloidal Superstructures. *Chem. Mater.* **27**, 8337–8344 (2015).

6. A. P. Thompson, *et al.*, LAMMPS - a flexible simulation tool for particle-based materials modeling at the atomic, meso, and continuum scales. *Comput. Phys. Commun.* **271**, 108171 (2022).

7. H. Jung, A. Yethiraj, A simulation method for the phase diagram of complex fluid mixtures. *J. Chem. Phys.* **148**, 244903 (2018).
